# Partial reprogramming restores youthful gene expression through transient suppression of cell identity

**DOI:** 10.1101/2021.05.21.444556

**Authors:** Antoine Roux, Chunlian Zhang, Jonathan Paw, José Zavala-Solorio, Twaritha Vijay, Ganesh Kolumam, Cynthia Kenyon, Jacob C. Kimmel

**Affiliations:** Calico Life Sciences LLC

**Keywords:** aging, reprogramming, cell identity, single cell RNA-seq

## Abstract

Transient induction of pluripotent reprogramming factors has been reported to reverse some features of aging in mammalian cells and tissues. However, the impact of transient reprogramming on somatic cell identity programs and the necessity of individual pluripotency factors remain unknown. Here, we mapped trajectories of transient reprogramming in young and aged cells from multiple murine cell types using single cell transcriptomics to address these questions. We found that transient reprogramming restored youthful gene expression in adipogenic cells and mesenchymal stem cells but also temporarily suppressed somatic cell identity programs. We further screened Yamanaka Factor subsets and found that many combinations had an impact on aging gene expression and suppressed somatic identity, but that these effects were not tightly entangled. We also found that a transient reprogramming approach inspired by amphibian regeneration restored youthful gene expression in aged myogenic cells. Our results suggest that transient pluripotent reprogramming poses a neoplastic risk, but that restoration of youthful gene expression can be achieved with alternative strategies.

## Introduction

Almost all metazoans experience aging, a process of progressive decline in functionality and resilience that results in death [1]. Germline development is the only known biological process capable of reversing or avoiding the effects of aging, and the ability to reprogram cells to a pluripotent state demonstrated that the stages of this developmental process can be reversed in individual cells [2]. Seminal work further demonstrated that cells could be reprogrammed to an induced pluripotent stem cell (iPSC) state through induction of four transcription factors – *Sox2, Pou5f1* (Oct4), *Klf4*, and *Myc* (SOKM) [3, 4].

Evidence from several groups further suggests that pluripotent reprogramming also reverses features of aging. iPSCs generated from young and aged donors are largely indistinguishable, and this similarity persists after differentiation into multiple fates [5, 6, 7]. This erasure of aging features does not occur in direct reprogramming protocols [7, 8], suggesting that dedifferentiation during pluripotent reprogramming is intimately related to this phenomenon.

Surprisingly, transient induction of SOKM in mice using an inducible germline allele was reported to improve multiple physiological functions in aged animals and to extend lifespan in progeroid mice [9]. This transient reprogramming strategy does not generate iPSCs, but rather alters gene expression without inducing a pluripotent state in most cells. Similarly, transient induction of pluripotency transcription factors was reported to reduce transcriptional features of aging in multiple human and mouse cell types [10, 11, 12] and to improve regenerative capacity in the muscle and eye [10, 11, 13].

These striking results suggest that even transient induction of pluripotency-associated gene regulatory networks (GRNs) is sufficient to ameliorate features of aging. However, it remains unclear if these effects occur uniformly in all cells and at which stage of the reprogramming process features of aging are lost. It is also unknown if transient reprogramming suppresses the GRNs that establish somatic cell identity. Early reports have suggested that cell identity programs are unaffected by transient reprogramming, but this stands in contrast to observations from iPSC reprogramming experiments [14, 15, 16, 17] and the known neoplastic effects of *in vivo* reprogramming [18, 9]. Likewise, it has been suggested that some of the beneficial effects of transient reprogramming stem from the re-engagement of somatic cell identity programs, but there is currently little evidence to support this hypothesis.

The necessity or sufficiency of individual pluripotency factors to ameliorate features of aging is also unclear. The widely used SOKM factors were identified based on their ability to induce a pluripotent state. Given that transient reprogramming explicitly avoids this outcome, it is plausible that only a subset of the SOKM GRNs are necessary to reverse age-related changes. Recent experiments suggest that *Myc* is not necessary at a minimum [11]. It is also unknown if alternative reprogramming strategies, such as multipotent reprogramming approaches, might also restore youthful gene expression, or if these effects are restricted to the activation of the full pluripotency program.

To address these questions, we performed transient pluripotent reprogramming using the Yamanaka Factors in adipogenic cells and mesenchymal stem cells (MSCs) isolated from young and aged mice and profiled gene expression by single cell RNA-seq. We found that transient reprogramming restored youthful gene expression but transiently repressed somatic cell identity in both cell types. To identify which GRNs in the Yamanaka Factor set were responsible for these effects, we performed a pooled screen for all possible combinations of factors and found that no single factor was required. We also tested a transient multipotent reprogramming regime in myogenic cells and found that features of aging were partially rescued.

## Results

### Transient reprogramming restores youthful gene expression in aged adipogenic cells

To evaluate the effects of transient reprogramming using the full Yamanaka Factor set, we designed a polycistronic tetracycline-inducible SOKM lentiviral vector with a constitutive fluorescent reporter (LTV-Y4TF, Fig. 1A)[19]. We used LTV-Y4TF and a tetracycline transactivator lentivirus (LTV-Tet) to transduce primary adipogenic cells from young (2-4 months old) and aged (20-24 months old) C57Bl/6 mice *in vitro*. Following transduction, we performed a pulse/chase by adding doxycycline (Dox) to the cell culture media for 3 days and chasing for 3 days (Fig. 1B). We used a multiplicity of infection (MOI) sufficient to transduce *<* 50% of cells in each condition, such that a fraction of cells in each well are exposed to Dox but do not contain both the transgenes required for reprogramming. These cells therefore serve as an *in situ* control. This experimental scheme provides a shorter induction period and a longer chase period than previous reports [9, 10, 11]. Following the chase period, we sorted dual transgene-positive (Tg+) and single transgene or transgene-negative (Tg-) cells of each age by cytometry and profiled cellular transcriptomes by single cell RNA-seq (Methods).

**Figure 1:**
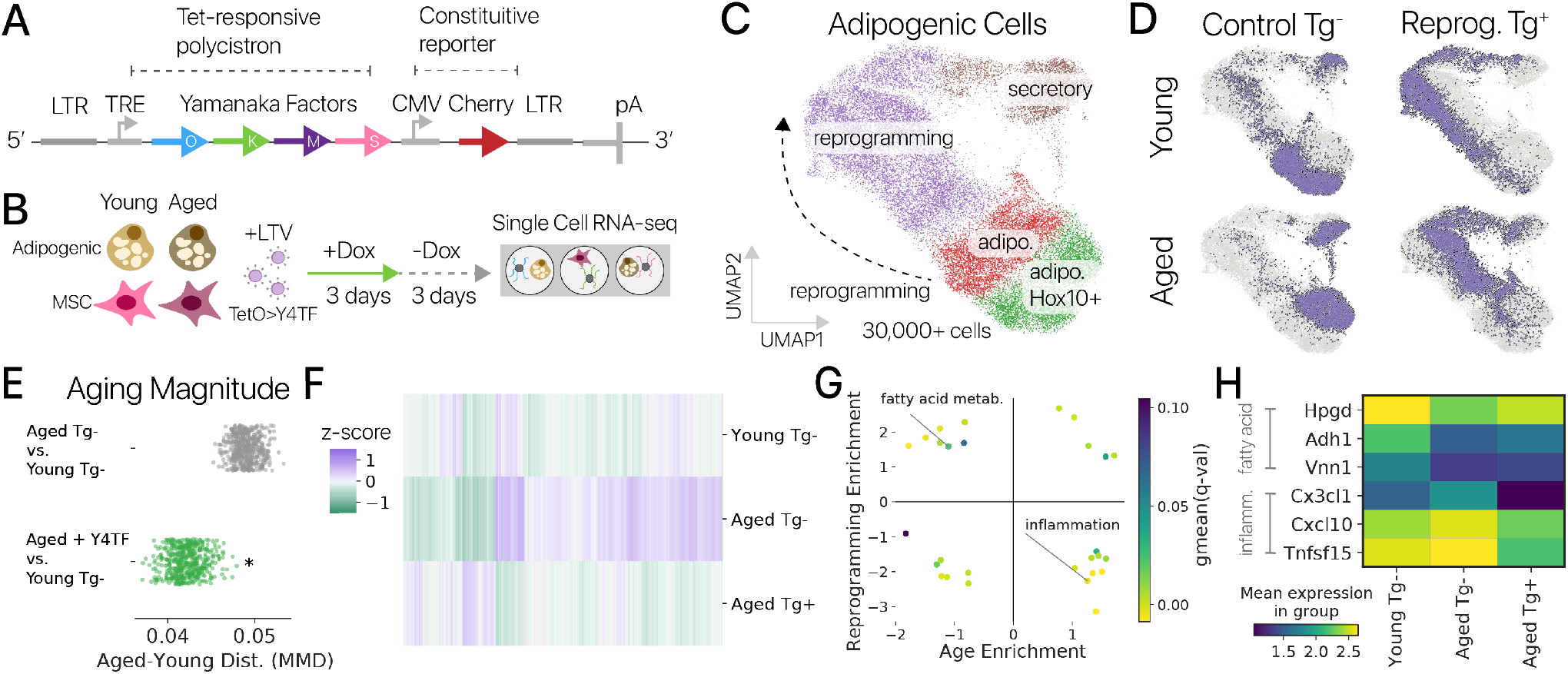
Transient pluripotent reprogramming restores youthful gene expression in murine adipogenic cells. **(A)** Schematic diagram of our pluripotent reprogramming lentiviral vector (LTV-Y4TF). Expression of a Yamanaka Factor polycistron is controlled by a tetracycline-response element (TRE). **(B)** Diagram of our transient pluripotent reprogramming experiment. We performed a 3 day pulse/3 day chase of SOKM using Dox inducer in adipogenic cells and muscle-derived MSCs from young and aged mice. After the chase, cells were profiled by single cell RNA-seq. **(C)** UMAP projection of single cell mRNA profiles from young and aged adipogenic cells. Control and reprogramming-specific expression states are annotated. Adipogenic states are marked by *Lpl*, secretory states by *Rspo1*, and reprogramming specific states by *Snca* and *Nanog*. **(D)** Control (Tg-) and reprogrammed (Tg+) populations of young and aged cells are highlighted in the embedding. Control adipogenic cells are divided into *Hoxc10*^*+/-*^ subsets. Young and aged populations are readily distinguished. **(E)** The magnitude of age-related change in gene expression was significantly decreased by Y4TF treatment, as measured using maximum mean discrepancy (MMD, *p <* 0.001, Wilcoxon Rank Sum). Aged reprogrammed cells are therefore closer to the young control state than aged control cells. **(F)** Transient reprogramming (Aged Tg+) restores youthful gene expression across thousands of genes (significant change in youthful direction, *q <* 0.10). **(G)** Gene set enrichment analysis shows that many gene programs downregulated with age are upregulated by reprogramming and vice-versa. Fatty acid metabolism was one of the youthful programs restored by reprogramming, and inflammatory responses upregulated with age were suppressed by reprogramming (MSigDB Hallmark gene sets). **(H)** Genes from the fatty acid metabolism and inflammatory response programs show amelioration of age-related change in transiently reprogrammed cells.

We captured 30,000+ adipogenic cell mRNA abundance profiles after quality control filtering. After denoising and dimensionality reduction with a variational autoencoder (Methods)[20], we clustered cell profiles and found that reprogramming induced a set of novel gene expression states unseen in control cells (Fig. 1C, D). Control cells occupied an adipogenic state marked by *Lpl* and a secretory state marked by *Rspo1*. The vast majority of treated cells occupied reprogramming-specific cell states, suggesting transient reprogramming effects are highly penetrant. Young and aged cells occupied distinct regions of the latent space in both control and reprogrammed populations, similar to a previous study of control cells *in vivo*, suggesting that some features of aging persist after reprogramming (Fig. 1D; Fig. S1)[21]. Animal-to-animal differences were a small source of variation (*<* 1%, Fig. S2, ANOVA, Methods).

We first wanted to determine if the transcriptional distance between young and aged control cells – the magnitude of aging – was decreased by transient reprogramming. We measured the magnitude of aging using the maximum mean discrepancy (MMD), a statistic from representation learning that measures the similarity of two populations and provides a test for statistical significance [22, 23]. Here, we computed the MMD across cell populations using autoencoder latent variables to capture changes across the entire transcriptome. We compared both aged control and aged reprogrammed cells to young control cells by MMD and found that the magnitude of aging decreased significantly after reprogramming treatment (Fig. 1E, *p <* 0.001, Wilcoxon Rank Sum; Methods). This decrease in the distinction of young and aged cells demonstrates that transient reprogramming can restore youthful gene expression across the transcriptome.

To determine which genes drive this change, we performed differential expression analysis and found that reprogramming treatment induced significant changes toward the youthful expression level in 3,485 genes out of a total of 5,984 genes changed with age (Fig. 1F). Reprogramming induced youthful changes significantly more often than would be expected by random chance (binomial test, *p <* 0.0001). We used gene set enrichment analysis [24] to compare the transcriptional effects of aging and reprogramming and found that reprogramming counter-acted age-related changes in many gene sets (Fig. 1G). Two of the strongest examples of this phenomenon were the adipogenic “fatty acid metabolism” gene set, downregulated with age and upregulated by reprogramming, and the “inflammatory response” gene set, upregulated with age and downregulated by reprogramming (Fig. 1H). Transient reprogramming may remodel the epigenome to allow for reactivation of the suppressed fatty acid program. We found a similar downregulation of adipogenic genes with age in single cell data collected *in vivo* (Fig. S1)[21]. Restoration of this characteristic metabolic program suggests that some functions of aged adipogenic cells may be improved by transient reprogramming.

### Cell identity dictates transient reprogramming effects

We next wondered if restoration of youthful gene expression was a consistent feature of transient reprogramming across cell types. We also performed transient reprogramming experiments in muscle-derived mesenchymal stem cells [25] to investigate. We captured 20,000+ MSC profiles and found that young and aged control cells were less distinct than in the adipogenic case (Fig. S3A, B), but cell age could still be readily distinguished by a classification model (93% accurate, Fig. S4, Methods). Similar to our experiments in adipogenic cells, transient reprogramming induced a novel set of reprogramming-specific gene expression states unseen in control cells of either age (Fig. S3C). These states were characterized by a downregulation of the epithelial-to-mesenchymal transition (EMT) program (Hallmark Gene Set enrichment of top 150 differential genes; Fischer’s exact test; *q <* 0.0001). We also found that youthful gene expression was restored in 712 genes (significant change in youthful direction), an aging-induced fibrotic gene program was suppressed, and an aging score derived from bulk RNA-seq data decreased (Fig. S3D, E; Fig. S5).

However, when we computed the magnitude of aging in MSCs, we found that aged cells were more distinct from young controls after reprogramming (Fig. S3F). This result was likely due to the fact that more than 4000 genes were significantly changed with reprogramming, but not with age. We hypothesized that the magnitude of aging might be smaller in MSCs than adipogenic cells, such that the orthogonal effects of reprogramming dominate in the MSC context. We tested this hypothesis by comparing the magnitude of aging in control adipogenic cells and MSCs in a shared transcriptional latent space and found that aging induced a significantly larger gene expression shift in adipogenic cells (Fig. S6A, C; Methods). These results suggest that transient reprogramming effects unrelated to cell age may dominate in some cell types.

We also hypothesized that reprogramming may induce some cell type specific effects. To test this hypothesis, we performed differential expression analysis to identify reprogramming-by-cell type interactions (Methods). This analysis revealed more than 10,000 genes for which reprogramming induced significantly different effects (*q <* 0.05, likelihood-ratio test, hurdle regression). We also found that 11% of non-residual variation was explained by cell type specific effects of reprogramming in a shared transcriptional latent space (Fig. S6B, ANOVA).

To unravel the influence of cell type on transient reprogramming outcomes, we extracted the top genes with significant cell type:reprogramming interactions and performed Gene Ontology analysis. Extracellular matrix and EMT programs were suppressed by reprogramming in both cell types, but significantly less so in adipogenic cells. By contrast, hypoxia and glycolysis gene sets were more upregulated by reprogramming in MSCs than adipogenic cells (Fig. S6E, *q <* 0.05, Fischer’s exact test). Investigating individual genes confirmed the influence of cell type on reprogramming effects in these gene programs (Fig. S6D, F). Somatic cell identity therefore dictates the effects of transient reprogramming across many genes, suggesting that youthful gene expression may be restored more effectively in some cell types than others.

### Somatic cell identity networks are repressed by transient reprogramming

Based on our observation of novel cell states in transiently reprogrammed adipogenic cells and MSCs, we hypothesized that the gene regulatory networks enforcing somatic cell identity may be repressed even though reprogramming factors were only expressed transiently. Given the continuum of reprogramming states we observed, a continuous coordinate system assigning each cell a position in the reprogramming trajectory enables parsimonious analysis. To build a coordinate system for transient reprogramming, we used pseudotime analysis, a method to infer continuous processes using the relationships among cells profiled at a single timepoint (Fig. 2A, C; Methods)[26].

**Figure 2:**
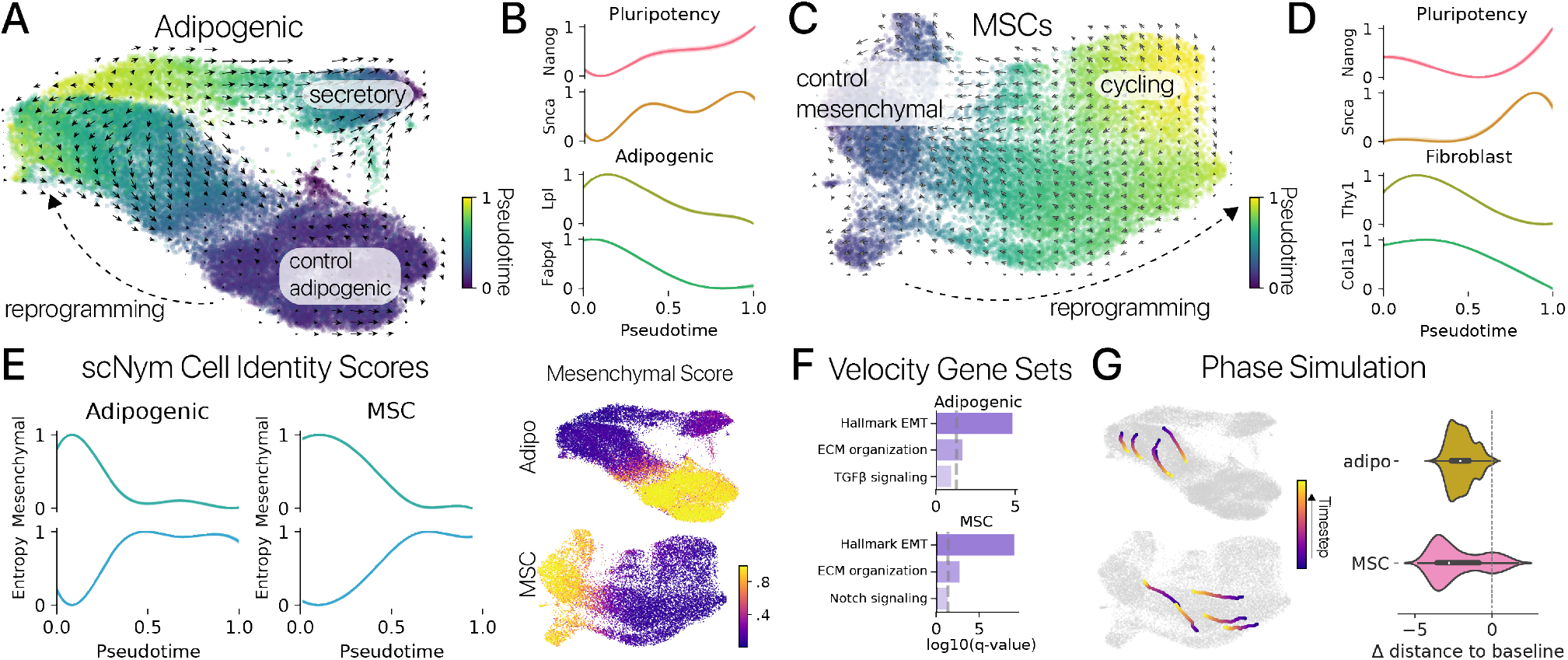
Somatic cell identity is suppressed by transient reprogramming and re-acquired by secondary differentiation. **(A)** Transient reprogramming in adipogenic cells induces a continuous trajectory of reprogramming-specific cell states. We captured this trajectory quantitatively using a diffusion pseudotime coordinate. RNA velocity (arrows) predicted that cells in reprogramming-specific states differentiate toward control cell identities. **(B)** Pluripotency associated genes *Nanog* and *Snca* were induced in the most distal reprogramming states, while adipocyte identity genes *Lpl* and *Fabp4* were suppressed (GAM fit values, mean *±* 95% CI). **(C)** Muscle-derived MSCs exhibit a similar trajectory of reprogramming states. RNA velocity similarly predicted that MSCs in reprogramming-specific states re-differentiate toward their original somatic cell identities. **(D)** Pluripotency associated genes were also induced in distal MSC reprogramming states, while mesenchymal identity genes *Thy1* and *Col1a1* are suppressed. **(E)** Mesenchymal identity program activity inferred using cell type classifiers and a mouse cell atlas dataset scored across pseudotime (left) and projected in transcriptional latent spaces (right). Mesenchymal identity programs were suppressed in distal reprogramming states for both cell types, while the entropy of cell identity increases (*p <* 0.01, Wald test). **(F)** Gene Ontology enrichment analysis reveals that epithelial-to-mesenchymal transition (EMT), extracellular matrix (ECM), and cell signaling genes are the major drivers of RNA velocity. **(G)** Cell state trajectories were simulated based on RNA velocity estimates in each cell type. Simulated cells were started in reprogramming states, and updated iteratively to predict their future expression. Simulated cells differentiated toward somatic cell states in both cell types (left). The distance between simulated reprogrammed cells and the control cell state decreased significantly (right) after 500 timesteps in the RNA velocity field (*n* = 1000 simulations; *p <* 0.01, one sample t-test).

We first investigated the effects of transient reprogramming on individual marker genes for somatic cell identity across the pseudotime trajectories. In adipogenic cells, we found that adipogenic genes *Lpl* and *Fabp4* were significantly downregulated in the most distal reprogrammed cells, while pluripotency associated genes *Nanog, Snca*, and *Fgf13* were upregulated (Fig. 2B, Methods). We found a similar pattern in MSCs where mesenchymal genes *Acta2, Thy1*, and *Col1a1* were downregulated while *Nanog, Snca* and *Fgf13* were upregulated (Fig. 2D). Activation of *Nanog* and other pluripotency genes after a 2-4 day SOKM induction is consistent with previous single cell timecourse studies of iPSC reprogramming (Fig. S7)[27]. Importantly, *Nanog* activation has also been proposed as the “point of no return” at which some cells may not regain somatic identities [28]. We next summarized the activity of cell identity GRNs using regulatory network inference methods [29, 30] and a cell identity classification model (scNym) trained on a mouse cell atlas and our data [31, 32]. We found that somatic cell identity programs were significantly suppressed in distal reprogramming states for both cell types using both analysis approaches (Fig. 2E; Fig. S8; *p <* 0.05, Wald test on cell identity predictions).

These results stand in contrast to previous reports that transient reprogramming did not suppress somatic cell identity or activate pluripotency genes [10, 11]. What could explain this difference? Previous studies relied on measuring a small set of marker genes in bulk transcriptional profiles. These assays may not have captured the suppression of cell identity or activation of pluripotency genes in a sub-population of cells. Our single cell profiling may therefore be capturing gene expression effects that are masked in bulk measurements. Supporting this view, our single cell measurements are consistent with lineage-tracing studies showing that transient pulses of SOKM induce *Nanog* expression and are sufficient to establish pluripotent colonies [15]. Our results suggest that transient reprogramming represses somatic cell identity GRNs and activates late-stage pluripotency GRNs in at least a subpopulation of cells in multiple cell types, potentially posing a neoplastic risk.

### Transiently reprogrammed cells re-acquire somatic identities through secondary differentiation

It has been proposed that transient expression of reprogramming factors induces a transient de-differentiation and re-differentiation process, such that reprogrammed cells temporarily adopt early phase reprogramming features but then reacquire their original state [33, 28, 10]. To date, there has been little evidence for this model in the context of restoring youthful gene expression. In addition to rich profiles of the current cell state, single cell RNA-seq provides information on future gene expression states through RNA velocity, allowing us to investigate the direction of cell state change in this trajectory [34]. RNA velocity infers a rate of change for each gene based on the relative ratio of spliced and unspliced transcripts under the assumption that unspliced transcripts represent nascent or newly transcribed messages.

We inferred RNA velocity for both our adipogenic cell and MSC experiments and found that velocity maps in both cell types suggest reprogrammed cells are re-differentiating toward their baseline state (Fig. 2A, C) [35]. To identify genes that drive predicted changes in cell state, we performed differential velocity analysis. In both cell types, we found that EMT and extracellular matrix gene sets were enriched in the velocity markers for reprogrammed cells, suggesting that transiently reprogrammed cells re-acquire their original mesenchymal identities (Fig. 2F).

We further quantified RNA velocity maps using phase simulations, a technique from dynamical systems to measure properties of vector fields by simulating the motion of a point moving through the field [36, 23, 37]. Here, we simulated reprogrammed cells in each velocity map and evolved their positions based on the inferred velocities of neighboring cells (Methods). We found that phase points initialized in reprogrammed states evolved toward control somatic cell states in both cell types (Fig. 2G). Paired with our previous result, our single cell analyses provide the first direct evidence that restoration of youthful gene expression involves the transient suppression of cell identity GRNs and subsequent re-differentiation and reactivation of these networks.

### Subsets of the Yamanaka Factors are sufficient to elicit transient reprogramming effects

Transient reprogramming studies in aged cells have largely investigated the effects of the canonical Yamanaka Factors (SOKM) along with additional pluripotency regulators known to induce pluripotency (*Nanog, Lin28*) [10, 4]. The pluripotency programs activated by the Yamanaka Factors contain positive feedback, so it is possible a subset of factors is sufficient to restore youthful expression, as suggested by a recent report using only SOK [11]. Several efforts have also demonstrated the sufficiency of Yamanaka factor subsets to generate iPSCs [38, 39, 40, 41], but it remains unknown if subsets exert similar transcriptional effects after only a short, transient induction. Both the full SOKM and reduced SOK factor combinations are oncogenic, motivating a search for alternative reprogramming strategies to restore youthful gene expression [18, 42].

Here, we investigated the effects of all possible sets of the Yamanaka Factors using a custom pooled screening system (Fig. 3A). Our system is inspired by other pooled screening and lineage tracing systems [43, 44, 45, 46] and encodes a tetracyclin-inducible reprogramming factor along with a constitutively expressed barcode on an individual lentivirus. We introduced these vectors in a pooled format in two independent experiments into young and aged MSCs and performed a 3 day pulse/3 day chase. The expressed lentiviral barcodes allowed us to demultiplex the unique combination of lentiviruses infecting each cell *in silico* (Fig. 3A, B; Methods). We confirmed the accuracy of demultiplexing by comparing to an orthogonal demultiplexing approach (Fig. S9, Methods).

**Figure 3:**
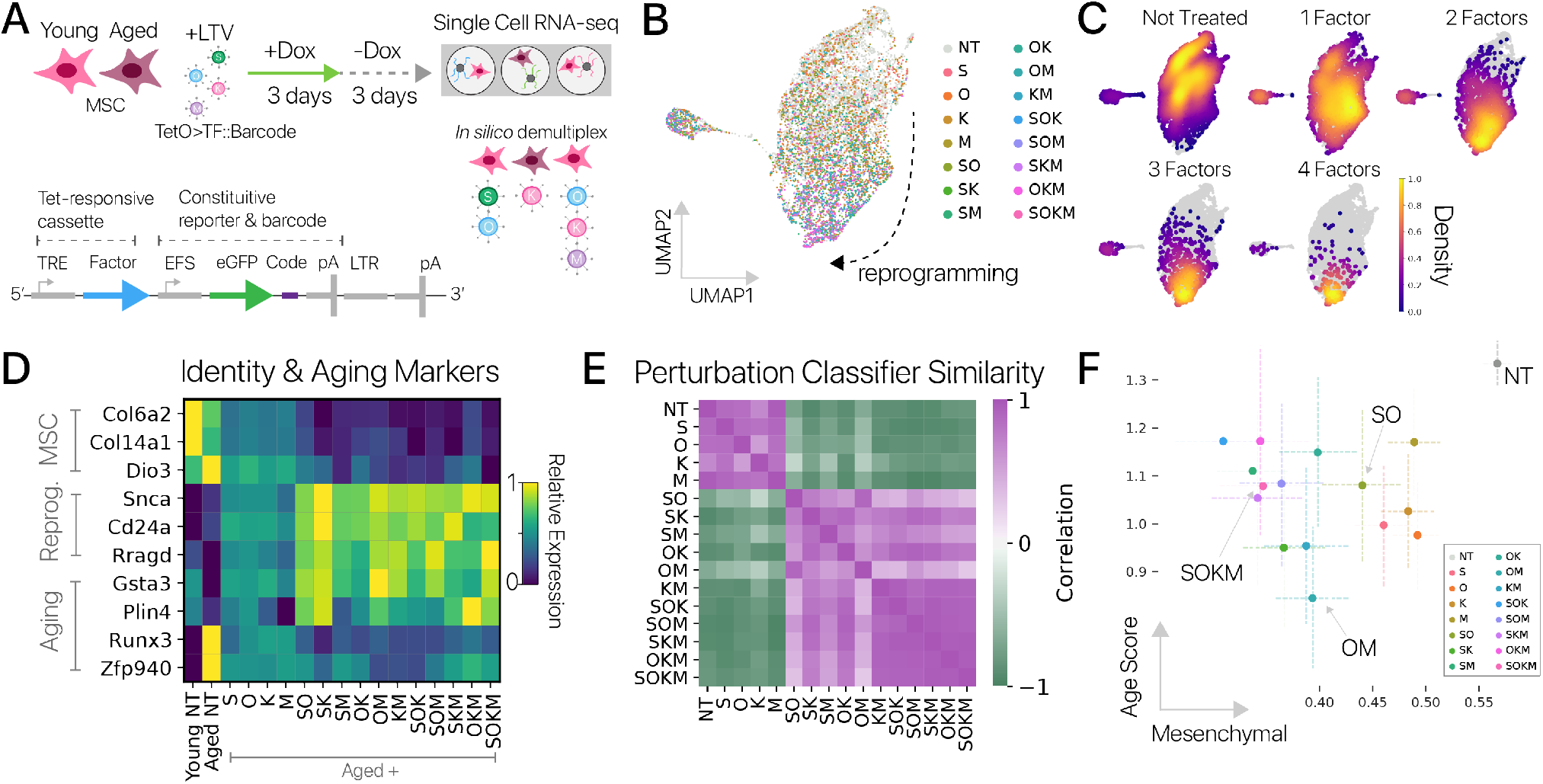
Pooled screening reveals the sufficiency of Yamanaka factor subsets and decoupling between rejuvenation and identity suppression. **(A)** Diagram of Yamanaka Factor pooled screening experiments. Young and aged MSCs were transduced with lentiviruses each harboring a one inducible Yamanaka Factor with expressed barcodes (lower). Reprogramming was induced for a 3 day pulse/3 day chase (*n* = 5 animals per age across two independent experiments). Cells were profiled by single cell RNA-seq and unique combinations of Yamanaka Factors were demultiplexed *in silico* based on expressed barcodes. **(B)** Cells from the pooled screen embedded using scNym, projected with UMAP, and labeled with the detected reprogramming factors (10,000+ cells). **(C)** Density of cells perturbed with different numbers of reprogramming factors in the UMAP embedding. Higher order combinations show a larger transcriptional shift relative to control cells. **(D)** MSC marker genes (top) decrease and reprogramming marker genes (center) increase as the combinatorial complexity (number of unique factors) increases. Aging genes (lower) likewise appear closer to the youthful level with more complex perturbations. **(E)** Similarity matrix between different Yamanaka Factor combinations extracted from a cell perturbation classification model. Similarity was computed as the correlation in prediction probabilities across cells for each pair of combinations. Low complexity perturbations and control (NT: not treated) cells are similar, while triplet combinations are highly similar to one another and the full Yamanaka Factor set (SOKM). **(F)** Mesenchymal cell identity scores derived from scNym models and aging gene set scores in aged cells reprogrammed with different factor combinations (mean scores and 95% CI). All but two Yamanaka Factor combinations significantly decreased both the mesenchymal cell identity score and age score relative to aged control cells (NT) (Wald tests, *p <* 0.01). However, age scores and identity scores were not well-correlated, suggesting that cell identity suppression and rejuvenation are not tightly coupled (Spearman *ρ* = *−*0.35, *p >* 0.20).

We recovered *>* 100 cells for all combinations of the Yamanaka Factors (Fig. 3B, C). To confirm the efficacy of our pooled screening system, we investigated the effects of each combination on reprogramming marker genes. We found that mesenchymal marker genes decreased as a function of combinatorial complexity (number of unique factors), while reprogramming marker genes increased (Fig. 3D). This result is consistent with the known biology of the Yamanaka Factors, where higher order combinations are more effective at reprogramming [3].

We next wanted to determine if different Yamanaka Factor combinations elicit unique effects, or if each combination induced similar gene expression programs, varying more in magnitude than direction. We investigated this question by training a state-of-the-art cell identity classification model to discriminate cells perturbed with each unique combination of Yamanaka Factors (Methods) [31]. Combinations that are highly similar might be frequently confused by the model, while combinations that are unique might be readily discriminated. To analyze our trained classifier, we extracted prediction probabilities across combinations for each cell and then computed the correlation of these probabilities for each pair of combinations as a similarity metric. We found that perturbations of similar combinatorial complexity were highly similar, while perturbations of different complexity were readily distinguished. For instance, most triplet combinations were frequently confused with the full Yamanaka Factor set (Fig. 3E). We also compared differentially expressed genes for each combination using the Jaccard index and mean expression vectors using the cosine similarity, yielding similar results (Fig. S10A, B).

Most combinations of Yamanaka Factors with similar complexity therefore have similar transcriptional effects when induced for a short period of time. The similarity among higher order combinations may be due to activation of shared effectors within the pluripotency network, even in the absence of missing factors, consistent with previous reports [38, 39, 40, 41]. It is also plausible that the endogeneous copy of the missing factor is activated by the remaining factor, as the Yamanaka Factors can activate one another [30](Fig. S10C). We did not observe significant up-regulation of missing factors in our data (*q >* 0.1, Monte Carlo differential expression), but we cannot rule out the possibility of transient activation. Taken together, our results suggest that no single Yamanaka Factor is required for restoration of youthful gene expression.

### Yamanaka factor subsets decouple rejuvenation and cell identity suppression

We wondered if suppression of cell identity and restoration of youthful gene expression were strongly associated across factor combinations, such that stronger restoration of youthful gene expression was predictive of a more suppressed identity program. To investigate, we scored a mesenchymal cell identity program using a cell identity classifier and the activity of an aging gene set extracted from our MSC experiments across cells in our screening experiment [31](Methods). We found that all combinations of Yamanaka Factors suppressed the cell identity program and reduced the aging score (Fig. 3). Aging score reduction was significant for all but two programs with lower cell numbers (SOK, OKM; Wald tests, *p <* 0.05).

However, suppression of the mesenchymal identity score and reduction in the aging score were not meaningfully correlated (Spearman *ρ* = *−*0.35, *p >* 0.20). This result was replicated when we considered each of the two independent experimental batches from our screen separately (minimum *q >* 0.12, maximum *ρ* = *−* 0.1 Spearman correlation). Some combinations reduced the aging score to roughly the same degree as the full Yamanaka Factor set, while suppressing mesenchymal identity significantly less (Wald test, *p <* 0.05). For example, the oncogene-free combination SO had a similar reduction in the aging score to SOKM, but suppressed mesenchymal cell identity 54% less than the full set. We repeated these experiments at a smaller scale in adipogenic cells and found similar qualitative results (Fig. S11). This result suggests that suppression of cell identity and restoration of youthful gene expression can be decoupled. It may therefore be possible to engineer reprogramming strategies that restore youthful expression while minimizing the risks of suppressed somatic identity.

### Transient multipotent reprogramming improves myogenesis in aged cells

Our results with the Yamanaka Factors suggest that alternative reprogramming strategies might also restore youthful expression. We wondered if transient reprogramming with multipotent reprogramming factors could also be effective. To test this hypothesis, we turned to a multipotent reprogramming system (as opposed to a pluripotent system) in myogenic cells inspired by limb regeneration in urodele amphibians using the transcription factor *Msx1. Msx1/2* are known to dedifferentiate myocytes to a multipotent state and have documented roles in digit and limb regeneration [47, 48, 49]. We performed a 3 day pulse/3-4 day chase of *Msx1* expression using a doxycycline-inducible lentiviral system in aged myogenic cells *in vitro*, then profiled cells by single cell RNA-seq in two independent experiments (Fig. 4A). As in our pluripotent reprogramming experiments, we sorted transgene positive and negative cells based on reporter expression to capture control and reprogrammed mRNA profiles (Fig. 4B, Methods).

**Figure 4:**
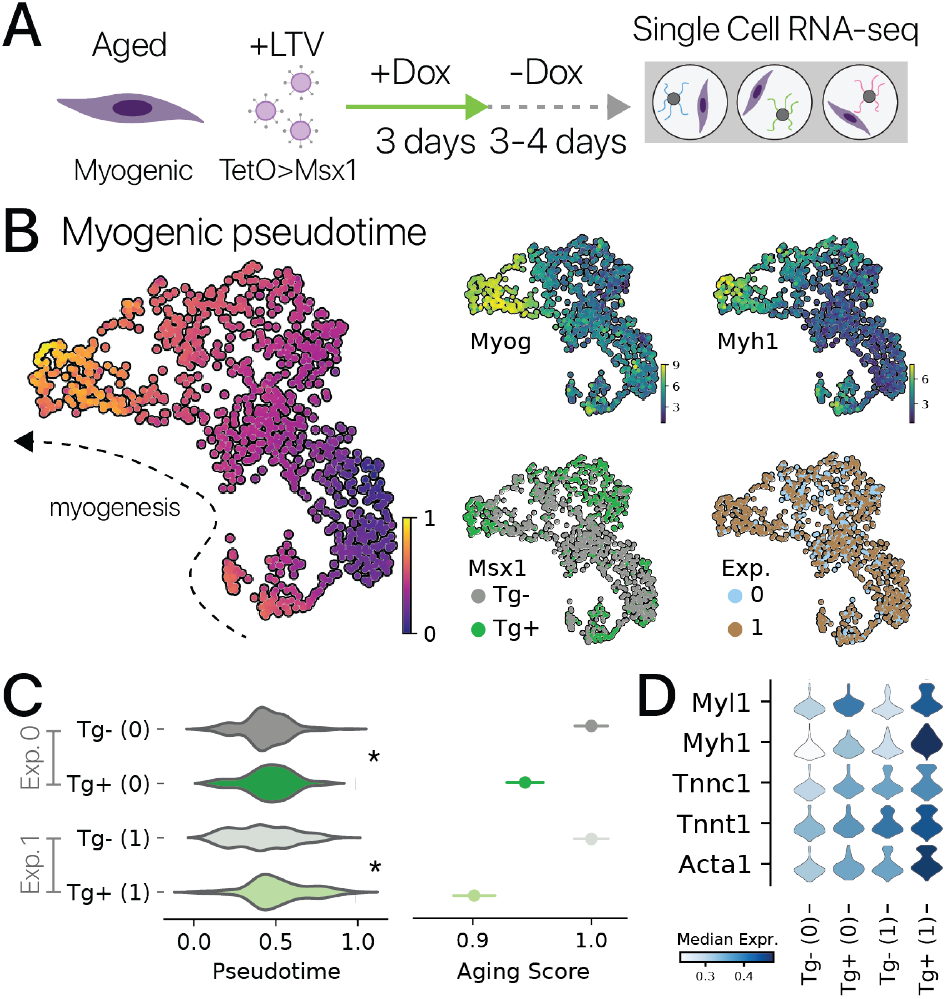
Pulsed *Msx1* reprogramming increases myogenic gene expression in aged myogenic cells. **(A)** Schematic of multipotent reprogramming experiments with aged myogenic cells isolated from (*n* = 2 and *n* = 3 animals). **(B)** Integrated cell profiles from two independent experiments (labels, lower right) projected with UMAP. Pseudotime analysis (left) provides a quantitative measure of myogenic differentiation state for each cell, confirmed by marker gene analysis (top right). **(C)** Transient multipotent reprogramming shifted cells into a significantly more differentiated state in both experiments (Wilcoxon Rank Sums, *p <* 0.05). Similarly, transient reprogramming significantly reduced a transcriptional aging signature extracted from a previous study [23] (mean *±* SEM; t-test, *p <* 0.01, normalized within experiments). **(D)** Myogenic marker genes for sarcomere components were enriched by transient multipotent reprogramming in both experiments (normalized within experiments, *q <* 0.1, Monte Carlo).

We captured myogenic cells in both progenitor (*Pax7+*) and differentiating states (*Myog+*). This is expected because our *in vitro* culture system provides a differentiation stimulus [50](Fig. 4B). It has previously been reported that aged myogenic cells do not differentiate into mature myocytes as effectively as young cells [51, 23]. We found that transiently reprogrammed aged cells were more differentiated than aged control cells based on pseudotemporal distributions, expression of marker genes, and gene set analysis (Fig. 4C, D; Fig. S12D). This result was robust to independent analysis of each experiment (Fig. S12A, B, C). We also measured an aging transcriptional signature derived from a previous study of myogenic differentiation and found that transient reprogramming significantly reduced the aging score (Fig. 4C), Methods, *p <* 0.01)[23]. The myogenic aging score itself is strongly influenced by the differentiation program, such that pseudotime explains a significant amount of variation in the aging score (*r*^2^ = 0.58, *p <* 0.0001 Wald test). These results suggest that multipotent reprogramming with *Msx1* can partially restore youthful gene expression in myogenic cells, similar to the Yamanaka Factors in adipocytes.

## Discussion

Aging induces broad gene expression changes across diverse mammalian cell types, and these changes have been linked to many of the prominent hallmarks of aging [52, 21, 53]. Cell reprogramming experiments have shown that young animals can develop from adult cells and aging features can be erased through complete reprogramming to pluripotency [2, 3, 5]. Recent reports have further suggested that transient expression of SOKM is sufficient to reverse features of aging and improve cell function [9, 10, 11, 12]. However, it was unclear whether these transient reprogramming interventions suppress somatic cell identities, activate late-stage pluripotency programs, or whether alternative reprogramming strategies could restore youthful gene expression.

Here, we investigated these questions using single cell measurements of gene expression to capture the phenotypic trajectory of transient reprogramming and evaluate the impact of alternative reprogramming methods. We found that transient reprogramming suppressed somatic cell identities and upregulated hallmark pluripotency programs (Fig. 2), contrary to some previous reports [10, 11] but consistent with timecourse iPSC reprogramming experiments (Fig. S7) and lineage-tracing studies of transient SOKM expression [15]. By inferring RNA velocity [34, 35] and applying numerical tools from dynamical systems [37], we also found that transiently reprogrammed cells transition back toward their original gene expression states after transit through an intermediate state (Fig. 2). Our single cell profiles therefore revealed transient cell states that were likely masked in previous bulk measurements and support a model in which transient reprogramming suppresses somatic identities that are later reacquired through differentiation. Further experiments profiling single cell populations at multiple time-points during transient reprogramming will be necessary to confirm this hypothesis.

It remains unknown which of the Yamanaka Factors are required to restore youthful gene expression, or which subsets might exhibit distinct effects during transient reprogramming. Previous studies have explored only one set of factors at a time, preventing accurate comparisons to address these questions [9, 10, 11, 12]. Our pooled screens of all possible Yamanaka Factor subsets revealed that combinations of 3-4 Yamanaka Factors have remarkably similar effects, suggesting no single factor is required to restore youthful gene expression. Combinations of two Yamanaka Factors were also more similar to the full SOKM set than to control or single factor perturbations, and all reprogramming factor combinations reduced an aging gene expression score (Fig. 3). Our screen demonstrates that no single pluripotency factor is required to mask features of aging and suggest oncogene-free reprogramming strategies may also restore youthful gene expression. Our multipotent reprogramming experiments in myogenic cells further support this suggestion, indicating that youthful gene expression may be restored even without activating the pluripotency factors (Fig. 4).

Restoring youthful gene expression can improve tissue function, implying that transient reprogramming may be therapeutic. However, pluripotent reprogramming is well-known to be an oncogenic process [18], even when *Myc* is excluded from the reprogramming set [42]. While it has been reported that transient reprogramming does not suppress somatic cell identities based on bulk measurements [9, 10, 11], our single cell results show that somatic cell identity is suppressed and late-stage pluripotency GRNs are activated in a transitional cell state in multiple cell types (Fig. 2). This raises the possibility that even transient reprogramming may be oncogenic. Identifying alternative reprogramming strategies to restore youthful gene expression with lower neoplastic risk is therefore desirable. Toward this aim, we have shown that transient reprogramming with multiple subsets of the Yamanaka Factors induces highly similar transcriptional effects to the full set, and that a distinct multipotent reprogramming system can confer youthful expression. These results suggest the feasibility of disentangling the rejuvenative and pluripotency inducing effects of transient reprogramming and serve as a resource for further interrogation of transient reprogramming effects in aged cells.

## Methods

### Animals

We isolated primary cells from the subcutaneous adipose tissue (inguinal pad) and limb muscles of young (2-4 months old) and aged (20-30 months old) C57Bl/6 male mice. Mice were ordered from the Jackson labs and aged at Calico Life Sciences, LLC. Young animals were allowed to acclimate for at least 3 weeks prior to experimentation. All animal experiments were approved by the Calico Institutional Animal Care and Use Committee, an AAALAC-accredited body.

We isolated cells from two animals of each age for adipocyte SOKM reprogramming experiments (aged animals, 28 months old) and three animals of each age for mesenchymal stem cell SOKM experiments (aged animals, 23 months old).We isolated MSCs from three animals of each age for one independent MSC pooled screening experiment, and from two animals of each age in a second independent experiment (aged animals, 28 months old). We isolated adipogenic cells from two animals of each age for one adipogenic pooled screening experiment (aged animals, 28 months old), and from three animals of each age in a second experiment (aged animals, 30 months old). For myogenic cell experiments, we isolated cells from two aged animals (28 months old) for the first experiment and three aged animals (30 months old) for the second experiment.

### Cell Isolation

We isolated pre-adipocyte cells from subcutaneous adipose tissue by dissecting tissue into small (1-2 mm) pieces with scissors and performing a 0.1% (w/v) collagenase II (Gibco) digestion in DMEM at 37^*o*^C for 1 hour. Adipose cell suspensions with washed in complete media (10% FBS, DMEM) and incubated in a 6-well plate overnight with one well per animal of origin.

We isolated mesenchymal stem cells from limb muscle tissue by dissecting muscle into small pieces with scissors and performing two enzymatic digestions. We first digested in 0.2% (w/v) collagenase II in DMEM for 90 minutes, then washed cell suspensions in digestion wash buffer (F10, 10% horse serum) and performed a second incubation in 0.4% (w/v) dispase II (Gibco) for one hour. Digested muscle cell suspensions were filtered through a 70 *µ*m cell strainer, then a 40 *µ*m cell strainer and stained with fluorophore-conjugated antibodies (see Table S1). We sorted mesenchymal stem cells as CD31^*−*^, CD45^*−*^, PI^*−*^, Sca1^+^ by FACS. MSCs were plated in one well each of a 6-well plate for each animal and incubated for 24 hours.

We isolated myogenic cells by performing the same digestion described for MSCs on limb muscle tissue. Once single cell suspensions were obtained, we performed a pre-plating procedure by incubating cells in a sarcoma-derived ECM coated well plate for 10 minutes, then immediately transferring cell suspensions to new ECM coated wells. This pre-plating step captures the majority of adherent, non-myogenic cells in the suspension and allows for capture of a high purity myogenic cell population in the final wells. Cells were incubated overnight in myogenic growth media (F10, 20% FBS, 1% Pen/Strep; Gibco, supplemented with 5 ng/mL rFGF2; R&D Systems).

### Cell culture

Adipogenic cells were passaged prior to transduction with lentiviral vectors. Cells were dissociated with TrypLE reagent (Gibco) and seeded at 100,000 cells/well in a 6 well plate, using separate wells for each animal and transduced after overnight incubation. Muscle-derived MSCs were passaged once prior to transduction using TrypLE. MSCs were seeded at 50,000 cells/well in a 6 well plate for polycistronic reprogramming experiments and 25,000 cells/well for pooled screening experiments. For arrayed screening experiments, MSCs were seeded at 7,500 cells/well in a 24 well plate, one plate per animal. For pooled screening in adipogenic cells, cells were seeded at 100,000 cells/well in 6 well plates. Myogenic cells were expanded for 5-7 days prior to lentiviral transduction in myogenic growth media and were passaged for seeding at 100,000 cells/well in 6 well plates for transduction. Myogenic cells were passaged with Cell Dissociation Buffer (Gibco).

### Lentiviral cloning and production

For polycistronic reprogramming experiments, we synthesized murine cDNAs for *Pou5f1, Klf4, Myc*, and *Sox2*. We removed stop codons from all cDNAs except *Sox2* and concatenated cDNAs into a polycistron using 2A-peptide sequences for a final open reading frame O-P2A-K-T2A-M-E2A-S. We inserted this OKMS polycistron into a 3rd generation lentiviral transfer vector flanked by a 5’ TRE3G tetracycline-inducible promoter and a 3’ woodchuck promoter response element. We also included a cytomeglovirus (CMV) promoter driven mCherry reporter transcript 3’ from the WPRE prior to the 5’ lentiviral terminal repeat (LTR). We refer to this vector as LTV-Y4TF.

For pooled screening experiments, we synthesized cDNAs for the Yamanaka Factors followed by an EF-1 alpha short (EFS) promoter driven eGFP with a unique 8-mer barcode in the 3’ UTR, followed by an internal SV40 polyA signal within the lentiviral genome. We cloned each Yamanaka Factor transgene into lentiviral transfer vectors with a 5’ TRE3G promoter to drive expression of the Yamanaka Factor. These designs were inspired by the CellTag lineage-tracing system [46] and allow us to detect the complement of Yamanaka Factors that transduced a given cell based on recovery of unique barcodes, even though the Yamanaka Factor transgenes are only transiently expressed. We refer to these vectors as LTV-S-BC, LTV-O-BC, LTV-K-BC, and LTV-M-BC where BC indicates a 3’ expressed barcode.

For multipotent reprogramming experiments in myogenic cells, we cloned the *Msx1* cDNA with a separable mCherry tag *Msx1-T2A-mCherry* into TRE3G driven lentiviral vector with a 3’ CMV driven eGFP reporter. We refer to this vector as LTV-Msx1.

We used three different tetracycline-transactivator lentiviral vectors, all harboring the same Tet3G tetracycline transactivator allele flanked by different fluorescent reporters to allow compatibility with different reprogramming vectors. For polycistronic reprogramming in muscle MSCs, we used a CMV driven Tet3G-T2A-mCerulean vector (LTV-Tet3G-mCerulean; adapted from VectorBuilder cat. VB180123-1018bxq by Gibson assembly). For polycistronic reprogramming in adipogenic cells, we used a vector harboring an EF1a driven Tet3G ORF followed by a CMV driven eGFP-T2A-PuromycinR reporter (LTV-Tet3G-eGFP; VectorBuilder cat. VB900088-2774nkq). For pooled screening, we used a vector harboring a CMV driven Tet3G followed by a 3’ WPRE and CMV driven mCherry reporter (LTV-Tet3G-mCherry; VectorBuilder cat. VB900088-2776tfj).

We packaged lentivirus by transfecting HEK293T cells with LV-MAX lentiviral packaging plasmids (Gibco) and the LTR-containing transfer vector of interest using Lipofectamine 3000 (Gibco). We collected supernatant from the viral packaging cells 24 and 48 hours after transfection, cleared supernatant by centrifugation, filtered supernatant with 0.45 *µ*m filters, and concentrated the cleared supernatant using PEG-it precipitation reagent (System-Bio). Lentiviral constructs were titered by transducing HEK293T cells with serial dilutions of virus and measuring fluorescent reporter expression frequency after 72 hours. We also prepared lentiviral particles through commercial vendors that use similar protocols.

### Lentiviral transduction

We performed lentiviral transductions using a standard spinfection method for all cell types. For adipogenic cells, we mixed the appropriate viral titer with complete growth media supplemented with [8 *µ*g/mL] polybrene and replaced growth media with the viral suspension. We centrifuged cells in well plates at 2000 x *g* for 1 hour and incubated overnight before exchanging growth media. For MSCs, we similarly added viral suspension, centrifuged cells at 2000 x *g* for 1 hour, and incubated cells overnight before exchanging media. For myogenic cells, we added viral suspension and transduced by spinfection for one hour at 1,500 x *g* with a polybrene adjuvant ([8 *µ*g/mL] in media).

### Transient polycistronic reprogramming

We performed transient reprogramming in adipogenic cells using LTV-Y4TF and LTV-Tet3G-eGFP. We transduced at MOI 30 in growth media containing polybrene ([8 *µ*g/mL]) by spinfection using a 1 hour centrifugation at 2000 *× g*. We replaced growth media after an overnight incubation and began a three day pulse of Dox ([4 *µ*g/mL]), exchanging media every 24 hours, followed by a three day chase.

We performed the same protocol for muscle-derived MSCs using LTV-Y4Tf and LTV-Tet3G-mCerulean each at MOI 10. For muscle-derived MSCs, we also included a control group transduced with virus, but never exposed to Dox. Following the chase period, cells were dissociated and we sorted cells expressing both LTV reporters (LTV-Y4TF : mCherry, LTV-Tet3G : eGFP or mCerulean) against all other cells by FACS. This sorting allowed us to enrich for transduced cells and profile treated (dual transgene positive) and untreated (missing at least one transgene) conditions to individual cells from within the same well, serving as an *in situ* control.

### Screening Yamanaka factor subsets

We performed a screen of Yamanaka factor subsets in two distinct experiments in MSCs. In the first experiment, we seeded cells in 24 well plates and delivered each combination of Yamanaka factors to one well of the plate at high MOI (MOI = 8) for each virus. We delivered a tetracycline-transactivator virus (LTV-*CMV>Tet3G-CMV>mCherry*) to all wells (MOI = 8). Before sequencing, we labeled the complexity of each perturbation (e.g. number of unique factors) with a unique cholesterol modified oligo to compare to our *in silico* demultiplexing. In the second experiment, we seeded cells in 6 well plates and delivered all Yamanaka factors in a pool to each well. We transduced one well per animal at both low and moderate MOIs (MOI = 3, 6) and pooled cells prior to sequencing. Two MOIs were used to increase representation of more complex perturbations. In both experiments, we pulsed Yamanaka factors by introducing doxycycline [4 *µ*g/mL] for three days and chased for 3 days. We likewise performed pooled screening in two distinct experiments for adipogenic cells. In the first experiment, we isolated cells from two young and two aged animals and transduced with MOI 8. In the second, we isolated cells from three young and three aged (30 months old) animals and trasduced with MOI 8. Adipogenic screens were performed with 100,000 cells/well in 6 well format.

### Transient multipotent reprogramming in myogenic cells

We performed transient reprogramming with *Msx1* in myogenic cells in two independent experiments. In both experiments, we seeded 100,000 myogenic cells/well in a 6 well plate in myogenic growth media. We transduced cells at MOI 15 with LTV-Msx1 and LTV-Tet3G-mCherry. We incubated cells for 12 hours then exchanged viral suspension with fresh growth media containing [4 *µ*g/mL] Dox. We replaced media daily for three days with Dox, then began a chase period where Dox-free media was used. We used a three day chase for the first experiment and a four day chase for the second. We sorted for cells expressing fluorescent reporters for both transgenes against other cells by FACS. This sorting allowed us to enrich for transduced cells and profile treated (dual transgene positive) and untreated (missing at least one transgene) conditions to individual cells from within the same well, serving as an *in situ* control.

### Single cell RNA-seq

We performed cell hashing with cholesterol-modified oligos (CMOs) to label cells from individual animals with unique barcodes. We used a unique group of of 1-2 CMOs (Integrated DNA Technologies) for cells from each animal and labeled cells following the MULTI-seq protocol [54]. Single cell RNA-seq libraries were prepared using the NextGEM v3.1 Single Cell Gene Expression chemistry (10x Genomics, Pleasanton, CA). Cells were emulsified using the Chromium controller (10x Genomics) and libraries were subsequently prepared following the library kit protocol. We ran young and aged cells in separate lanes of the 10x instrument for all experiments to provide the highest fidelity sample demultiplexing. For all experiments except the pooled screens, we additionally ran transgene-positive cells carrying both the reprogramming vector and a tetracycline-transactivator and transgene-negative cells (lacking at least one vector) in separate lanes. We added the MULTI-seq additive primer during the cDNA amplification step to allow for CMO barcode amplification, as described in the MULTI-seq protocol. For myogenic reprogramming experiments, we combined myogenic cells with highly distinct cell types prepared for unrelated experiments in the same lane and extracted myogenic cells *in silico* for analysis. We also pooled myogenic cells derived from different animals without CMO barcoding to avoid cell loss for this rare cell type.

### Read alignment and cellular demultiplexing

We pseudoaligned reads to the mm10 reference genome using “kallisto” and performed cell barcode demultiplexing and UMI read aggregation using the “kallisto | bus-tools” workflow [55, 56]. For experiments with transgenic constructs, we modified the mm10 reference to include lentiviral genomes as additional chromosomes in the reference genome. This allowed us to detect the presence of transgenic transcripts in our sequencing data.

We assigned MULTI-seq library reads to cell barcodes using “kite” [57]. For pooled screening experiments, we additionally quantified the number of lentiviral transgene barcodes in each cell using “kite”.

### mRNA profile denoising, latent variable inference, and data integration

We denoised mRNA abundance profiles using scVI [20] models with 64 latent variables. We fit scVI models for 1000 epochs using early stopping to minimize the evidence lower-bound (ELBO). For MSC pooled screening experiments, we integrated our arrayed and pooled transduction experiments by injecting batch covariates into the scVI model. Similarly, we used batch covariate injection to integrate across independent adipogenic pooled screening experiments and myogenic multipotent reprogramming experiments. For the latter application, we reduced the latent space size to 32 variables to avoid overfitting to a smaller dataset. For myogenic experiments, we additionally employed “harmony” integration on a PCA decomposition of the scVI latent space to account for batch effects across independent experiments [58]. To construct a shared latent space between adipogenic cells and MSCs, we fit an scVI model to both cell populations simultaneously. For all experiments, we constructed a nearest neighbor graph in the scVI latent space and projected this graph into two dimensions for visualization with UMAP.

### MULTI-seq demultiplexing

We used the “hashsolo” approach to demultiplex MULTI-seq read counts in both adipogenic and MSC experiments with a prior distribution of [0.02, 0.94, 0.04] for negative, singlets, and doublets respectively [59]. We manually verified “hashsolo” classifications by clustering cell profiles using CMO counts. We adjusted cluster labels based on manual inspection where appropriate. We only used confident classification calls for animal-specific analysis.

### Bulk RNA-seq of young and aged MSCs

We performed bulk RNA-seq on young and aged MSCs in three independent experiments. Cells were collected from three young (3-4 months) and three aged (20-24 months) animals in each experiment. In the first and second experiment, freshly-isolated cells were cultured for 11-14 days before RNA collection In the third experiment, cells from the second fresh cell experiment were cryopreserved in Recovery Cell Culture Freezing Media (Gibco), then thawed and cultured for 7 days before RNA collection. RNA was isolated from all cell using a Zymo Quick RNA kit (Zymo Research) and RNA-seq libraries were prepared with NEB-Next Ultra II Directional RNA Library Prep Kit (New England Biolabs). Libraries were sequenced on an Illumina NovaSeq. We pseudoaligned reads with kallisto [55] and quantified differential expression with sleuth [60]. We extracted an aging gene signature by selecting genes that changed with age under a loose FDR threshold (*q <* 0.4) across both freshly isolated cell experiments and cryogenically preserved cell experiments. This procedure yielded a signature of 198 aging genes which we used to score aging gene expression in our MSC SOKM reprogramming experiment.

### Estimating the contribution of covariates to total variation

We estimated the contribution of experimental covariates (age, transgene treatment) to the total variation observed in our experiments using ANOVA. We treated the scVI latent encoding as a set of response variables and fit linear models for each variable of the form *z*_*j*_ *∼*age + treatment + age:treatment where *z*_*j*_ is a latent variable. We also assessed the contribution of the animal-of-origin for each cell using animal-specific labels derived from MULTI-seq demultiplexing. We performed ANOVA as above with animal-specific labels using models of the form *z*_*j*_ *∼* age*treatment + animal, where “animal” is a label specific to each “age:treatment” combination because we ran “age:treatment” combinations in separate library preparation reactions. We included “negative” classifications from “hashsolo” in this regression in case a CMO hashing failure captured structured variation.

### Differential expression

We performed differential expression testing across binary contrasts by estimating log-fold change distributions for each gene using Monte Carlo approximations from the scVI posterior distribution [61]. We estimated the false discovery rate (FDR) of differential expression as the fraction of Monte Carlo samples that did not show a log-fold change above a minimum threshold (|log_2_ *a/b* |*≥*0.5). To test continuous covariates and cell type:reprogramming interaction effects, we used logistic/Gaussian hurdle models inspired by MAST [62] for individual genes and logistic models for gene program scores on the unit interval [0, 1], as previously described [23]. We performed FDR control for MAST models with the Benjamini-Hochberg procedure [63]. We performed differential expression testing on pseudotime covariates using models of the form “Gene *∼*Pseudotime”. We tested cell type:reprogramming interactions with models of the form “Gene *∼*Age + Cell Type + Treatment + Cell Type:Treatment”.

### Gene set enrichment analysis

We performed enrichment analysis for Gene Ontology terms using Enrichr [64]. We performed rank-based Gene Set Enrichment Analysis (GSEA) using the “fast GSEA” implementation of the GSEA algorithm [24] and gene sets extracted from MSigDB [65].

### Generalized additive modeling of gene expression over pseudotime

We fit generalized additive models (GAMs) to express relationships between genes or gene programs and continuous pseudotime coordinates. We fit Gaussian GAMs of the form “Gene *∼* Pseudotime” using six cubic splines through the PyGAM framework [66]. We represent expression trends over pseudotime using the mean value of GAM predictions along with the 95% confidence interval.

### Estimating aging magnitudes in transcriptional space

We estimated an aging magnitude as the difference between young and aged cell populations. We estimated this difference between two populations using the maximum mean discrepancy (MMD) computed on scVI latent variables. scVI latent variables capture an expressive, low-dimensional representation of gene expression and are therefore useful for this task. We computed the MMD our previously described “scmmd” package over a series of bootstrap random samples to ensure robustness [23]. Briefly, we sampled *n* = 300 cells from each population and computed an MMD at each of 500 iterations. We performed these MMD comparisons to compare young, untreated cells to both aged, untreated cells and aged, reprogrammed cells. We assessed the significance of changes in the MMD using the Wilcoxon Rank Sum test across bootstrap iterations.

### RNA velocity analysis

We counted estimated the fractions of spliced and un-spliced reads for each transcript based on the alignment of reads to exons and introns using “kallisto | bustools” to an mm10 reference genome from Ensembl [55, 56]. We estimated RNA velocity [34] using the stochastic model implemented in “scvelo” [35].

### Pseudotemporal trajectory inference

We inferred pseudotime trajectories for adipogenic cells and MSCs using “scvelo”. Briefly, we constructed a nearest neighbor graph in the embedding space and weighted edges in the graph based on the directionality of RNA velocity vectors such that neighbors in the path of the RNA velocity vector received higher transition probabilities. We then paired this matrix with the diffusion pseudotime inference method [67]. We inferred pseudotime trajectories for myogenic cells using diffusion pseudotime analysis on a nearest neighbor graph constructed in diffusion component space. We selected root cells from within the top decile of *Snai2* (primitive marker gene) expression.

### Phase simulations in RNA velocity fields

We performed phase simulations in RNA velocity fields using our previously described “velodyn” package [23]. For both adipogenic cells and MSCs, we initialized *n* = 1000 phase points within the transiently reprogrammed cell population at positions *x*_0_ and evolved their positions based on the RNA velocity vectors of their neighbors for *t* = 500 timesteps with a step size of *η* = 0.5. We used an update rule

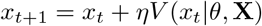

where we update the current position *x*_*t*_ to a new position *x*_*t*+1_ using an update function *V* (*x*_*t*_ |*θ*, **X**) that infers a velocity vector *v*_*t*_ from *k*-nearest neighbors given parameters (*k, η*). We parameterize *V* as a stochastic function sampling from a weighted multivariate Gaussian distribution 𝒩 (*µ*^*∗*^, Σ^*∗*^) where neighbors are weighted based on the distance in the embedding space from the query point.

### Cellular age classification

We trained semi-supervised variational autoencoder models using the scANVI framework to discriminate cell age [68]. Prior to model training, we split our data into a training set (80%) and held-out test set (20%) with stratification. We trained scANVI models for 100 unsupervised epochs (reconstruction loss only) and 300 semi-supervised epochs (reconstruction and classification losses) using 10% of the training set as validation data for early stopping. We evaluated model performance based on the accuracy of classifications in the held-out test set, unseen during model training.

### Embedding reprogramming factor perturbations with scNym

We fit scNym cell identity classification models to discriminate MSCs transiently reprogrammed with different Yamanaka Factor subsets [31]. We subsampled a balanced training set from the full MSC pooled screening data by extracting *n* = 100 cells per factor combination. We treated all remaining cells as a held-out test set. We trained an scNym model on the training set using *n*_hidden_ = 128 hidden units per layer, a patience period of 30 epochs prior to early stopping, a maximum number of 150 epochs, and default settings for all other parameters.

We subsequently predicted factor combinations for all cells in the MSC pooled screening experiments. We projected the scNym embedding activations for visualization using UMAP. We computed the correlation of scNym predictions for each factor combination by computing correlations among the columns of the scNym predicted class probability matrix *ŷ*^Cells*×*Combinations^.

### Scoring cell identity programs with scNym

We used scNym models to score the activity of specific cell identity programs in transiently reprogrammed cells. We fit semi-supervised, adversarial models using the *Tabula Muris* as a training set and our transiently reprogrammed cells as a target data set. We used the “cell ontology class” annotations in the *Tabula Muris* as class labels. We fit models for up to 200 epochs and used early stopping on a validation set held-out from the training data to select the best performing model. We fit separate models for the adipogenic SOKM reprogramming experiment, the MSC SOKM reprogramming experiment, and the MSC pooled screen.

We used trained models to both derive an embedding and predict cell identities from within the *Tabula Muris* training set in our transiently reprogrammed cells. We summed the likelihood of several similar cell identities to create a “Mesenchymal Identity” score for each cell (identities: “mesenchymal stem cell”, “mesenchymal cell”, “stromal cell”). We computed the entropy of cell identity as the Shannon entropy of the cell identity probability vector predicted by scNym.

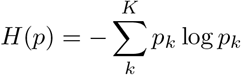

where *k ∈ K* are class indices, *p* is the identity probability vector, and *H*(*·*) is the entropy function.

### Scoring aging gene expression in pooled screening experiments

To compare the impact of different Yamanaka Factor combinations on features of aging, we devised an “Aging Score” based on genes differentially expressed with age in MSCs. We first identified genes that were significantly differentially expressed (*q <* 0.10) the same direction in control MSCs for both our polycistronic reprogramming experiment and the pooled screens. We took the intersection of this set with the set of genes that showed a significant restoration of youthful gene expression in any of the Yamanaka Factor combinations. This yielded a set of 228 aging genes that are influenced by reprogramming. We derived separate Age_Up_ and Age_Down_ scores using a standard gene set scoring method for the upregulated and downregulated genes respectively [69]. We then took the difference of these scores as the Aging Score: Aging Score = Age_Up_ *−*Age_Down_. We used a similar procedure for adipogenic pooled screening experiments to generate a set of 801 aging genes and an associated aging score.

To quantify the effect of each combination on the age score, we fit a linear regression of the form: Aging Score *∼*Combination + Age using aged cells for all treatments and only control (NT) young cells as observation data. We exclude young treated cells from the regression so that coefficients for each combination represent the effect of the combination on aged cells. We extracted coefficients and their confidence intervals to compare across combinations (Fig. 3F).

### Scoring aging gene expression in myogenic multipotent reprogramming experiments

We re-analyzed publically available single cell RNA-seq profiles of young (3 months old) and aged (20 months old) myogenic cells after *in vitro* differentiation to identify an aging gene signature (GEO : GSE145256)[23]. We extracted 329 genes that were changed more than 2-fold between young and aged myogenic cells with a false discovery rate *q <* 0.10 (Wilcoxon Rank Sum test, Benjamini-Hochberg FDR control). We then generated Age_Up_ and Age_Down_ scores as described above to construct an aging score. For visualization, we Winsorized the aging score to the [3, 97] percentile interval.

### Re-analysis of iPSC reprogramming timecourse

We re-analyzed data from a previous single cell RNA-seq study of mouse embryonic fibroblast to iPSC reprogramming (GEO : GSE122662)[27]. We trained an scVI model to denoise the data, learn latent variables, and perform differential expression as above.

### Re-analysis of Tabula Muris Senis adipose tissue aging

We re-analyzed data from the *Tabula Muris Senis* single cell RNA-study of mouse aging in the adipose tissue (GEO : GSE149590)[21]. We normalized expression data with a log(CountsPerMillion+1) transformation, selected highly variable genes, and embedded cells with UMAP on a PCA derived nearest neighbor graph. We performed differential expression with a two sample t-test using scanpy [70].

## Author’s Contributions

AR designed and performed molecular biology experiments and contributed intellectually. CZ and JZS performed animal experiments. JP performed cytometry experiments. TV performed bulk RNA-seq experiments. GK supervised research. CK contributed intellectually and edited the paper. JCK conceived the study, designed experiments, performed cell culture, molecular biology, reprogramming, and single cell RNA-seq experiments, analyzed data, supervised research, and wrote the paper.

## Supplemental Figures

**Figure S1:**
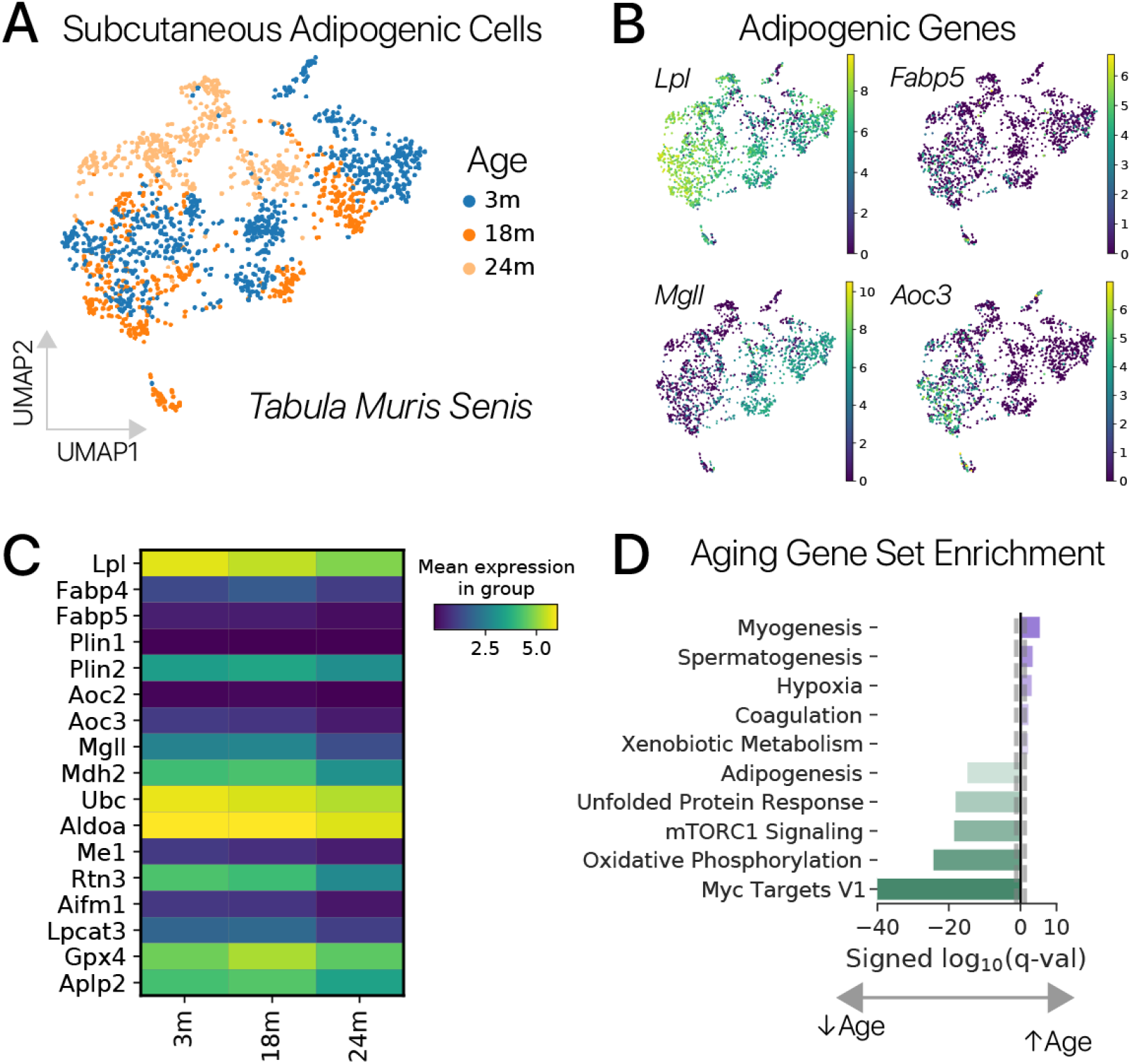
Adipogenic cells *in vivo* show similar features of aging to adipogenic cells *in vitro*. **(A)** Adipogenic cells from the subcutaneous adipose tissue in the *Tabula Muris Senis* [21] embedded with UMAP, colored by age. Young cells (3m, months) and aged cells (24m) clearly separate in transcriptional space, similar to our *in vitro* data. **(B)** Expression of several adipogenic genes is restricted to young cells. **(C)** Many adipogenic genes show significantly decreased expression with age, similar to *in vitro* results (*q <* 0.05, *t*-test). **(D)** Gene set enrichment analysis for genes significantly changed between 3 months of age (3m) and 24 months of age (24m) reveals strong downregulation of Adipogenesis, Oxidative Phosphorylation, and Unfolded Protein Response programs (Hallmark MSigDB, *q <* 0.01, Fisher’s exact test).

**Figure S2:**
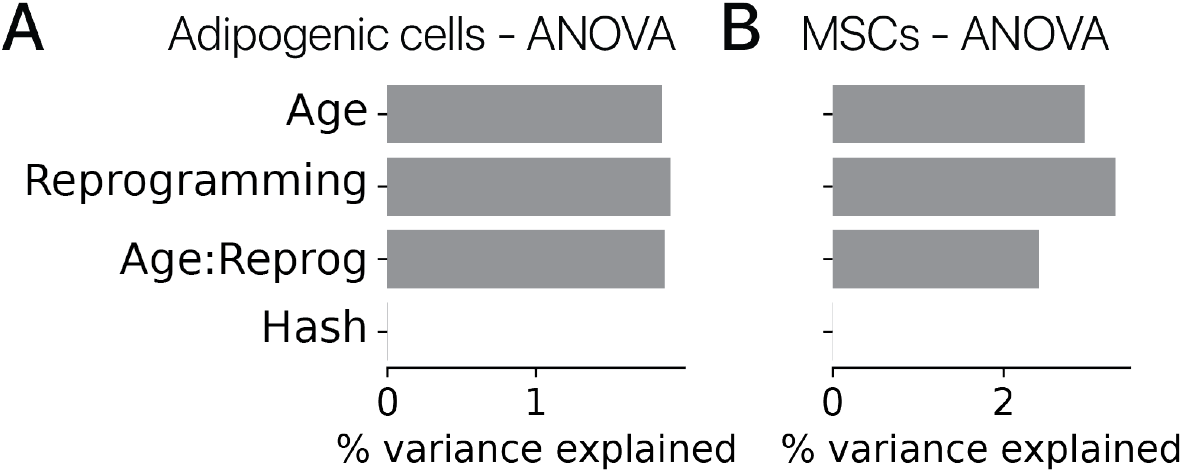
Age and reprogramming effects are the dominant sources of variation in transiently reprogrammed cells. **(A)** ANOVA results in transiently reprogrammed adipogenic cells, where the remaining variation is unexplained by the covariates shown. **(B)** ANOVA results in transiently reprogrammed MSCs. In both cell types, Age and Reprogramming Treatment are the major source of explained variation, while the animal of origin (Hash) is a minor contributor.

**Figure S3:**
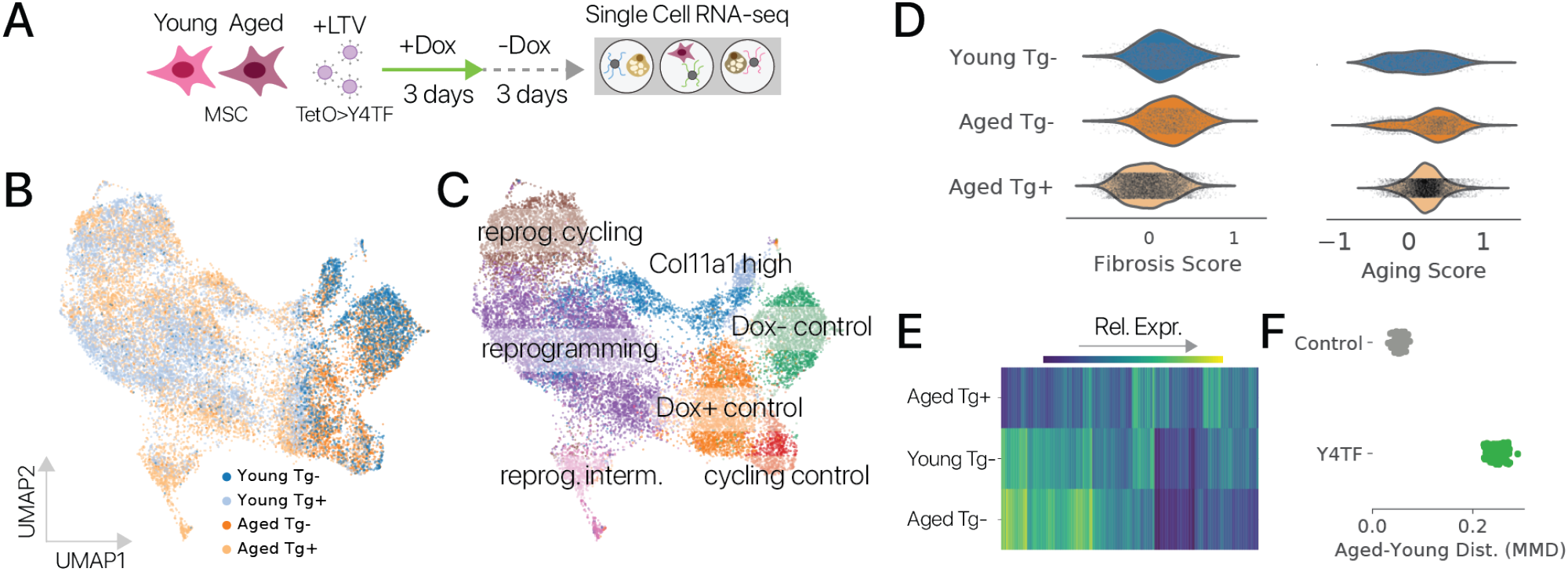
Transient reprogramming in muscle-derived MSCs remodels gene expression. **(A)** Experimental schematic. Muscle-derived MSCs were transduced with a tetracycline-inducible polycistronic Yamanaka Factor cassette and transiently reprogrammed with a 3 Dox day pulse, 3 day Dox chase. **(B)** Reprogrammed cells (Tg+) clearly segregate from control cells (Tg-) in a latent transcriptional space, while age is smaller source of variation. **(C)** Single cell profiles resolve distinct cell states induced by reprogramming. **(D)** Transient reprogramming ameliorates aging-induced increases in fibrosis gene sets and an aging gene score derived from bulk RNA-seq experiments. **(E)** Youthful gene expression is restored in thousands of genes by transient reprogramming, despite the large magnitude of reprogramming effects orthogonal to the axis of aging. **(F)** Transient reprogramming increases the maximum mean discrepancy between young control cells and aged cells, suggesting that many effects of reprogramming are orthogonal to the axis of aging.

**Figure S4:**
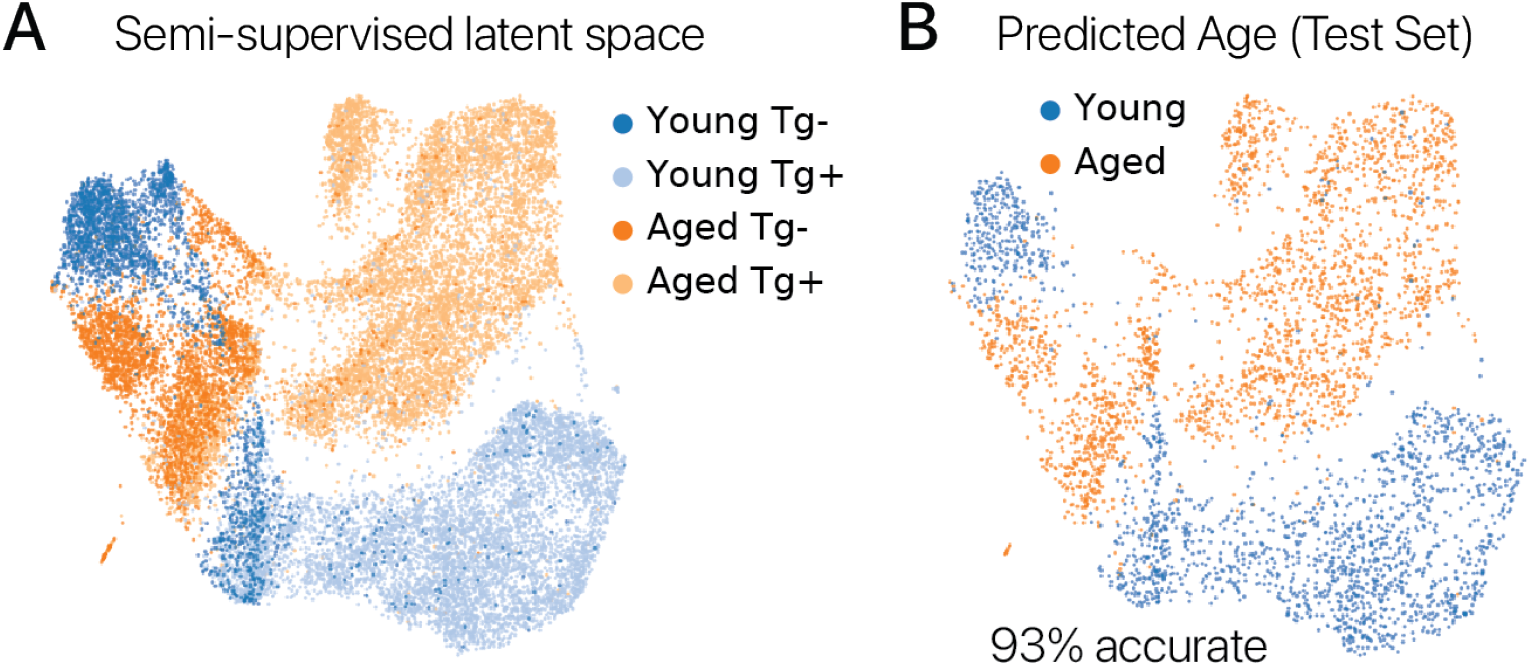
Semi-supervised aging classifiers demonstrate segregation of MSCs by age in transcriptional space. **(A)** MSCs embedded in a latent space using a semi-supervised variational autoencoder model. We trained variational autoencoder to jointly reconstruct transcriptomes and classify cell age, learning a representation that segregates young and aged cells in the latent space. **(B)** Cells from the test set projected in the semi-supervised latent space colored with the class predictions. We found that age predictions were 93% accurate, confirming that features of aging are captured at the transcriptional level.

**Figure S5:**
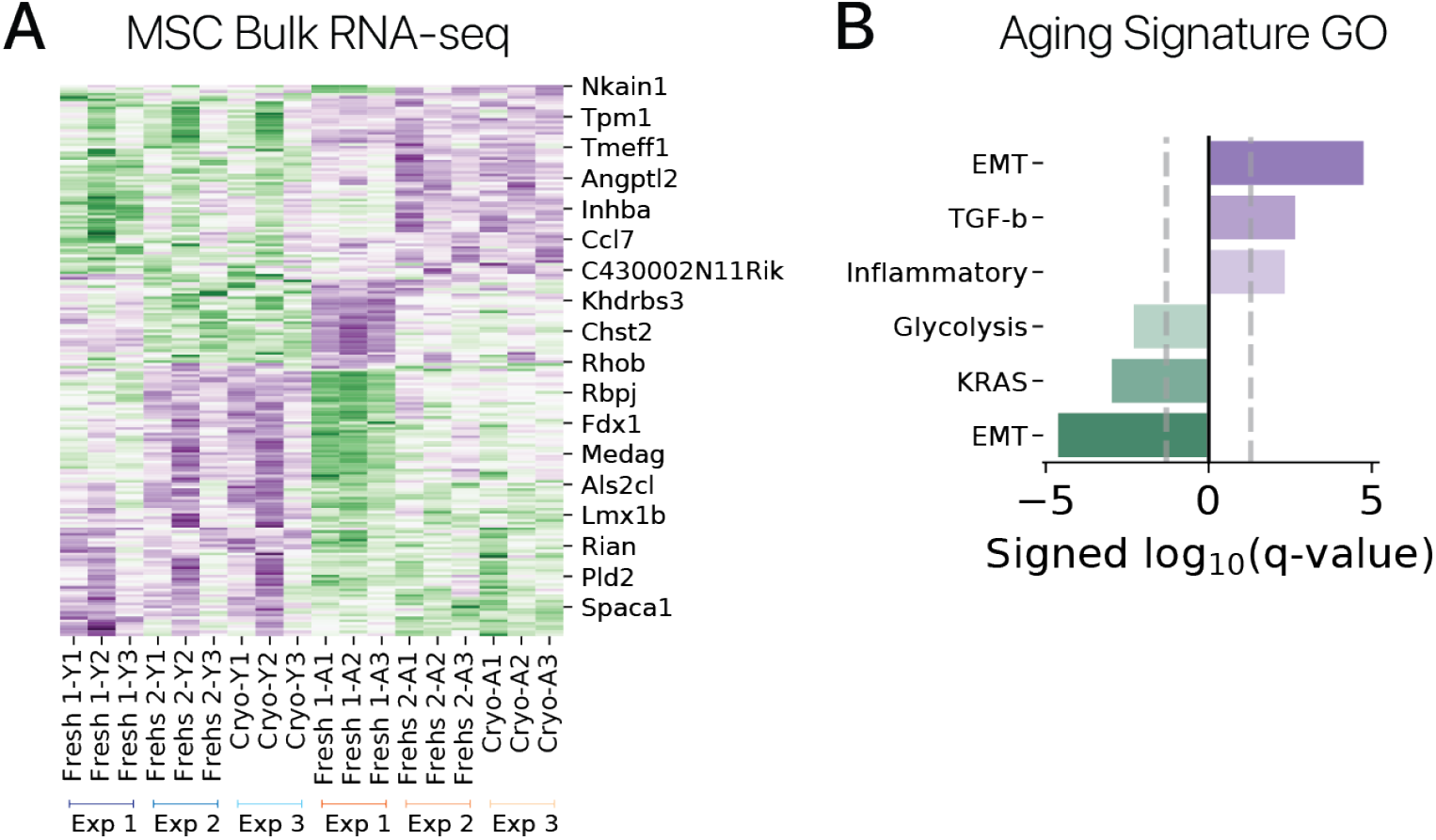
Bulk RNA-seq reveals a reproducible transcriptional signature of aging in MSCs. **(A)** Bulk RNA-seq profiles of young (Y) and aged (A) MSCs collected in three separate experiments. Genes with significant aging effects in all three experiments are shown. We used these gene sets to derive an “Aging Score” for our transient reprogramming experiment (Fig. S3D). “Fresh” samples were prepared from freshly isolated MSCs after an *in vitro* culture period, while “Cryo” samples were prepared after cryopreservation and subsequent culture. **(B)** Gene Ontology enrichment analysis for MSigDB Hallmark gene sets. Epithelial-to-mesenchymal transition (EMT) genes are strongly enriched for age-related changes. Both inflammatory genes and the fibrotic signaling factor TGF-*β* show age-related increases.

**Figure S6:**
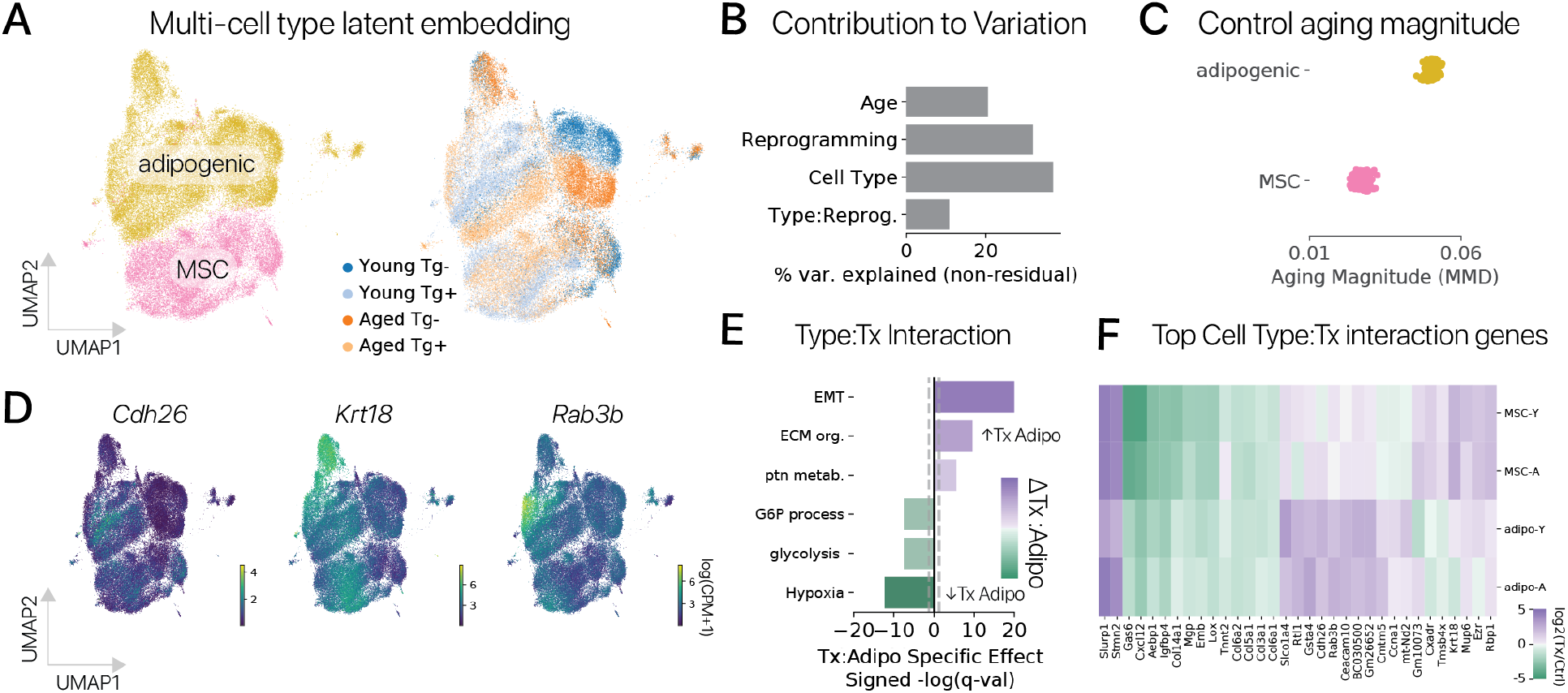
Cell identity dictates the effects of transient reprogramming. **(A)** Transiently reprogrammed adipogenic cells and MSCs from our polycistronic experiments were embedded in a joint latent space with scVI to allow for direct comparison of effect sizes. **(B)** ANOVA revealed that Cell Type:Reprogramming interactions account for roughly 11% of the total non-residual variation explained by covariates. **(C)** We re-computed aging magnitudes by maximum mean discrepancy for control adipogenic cells and MSCs in the joint latent space to allow for direct comparisons. We found that the magnitude of aging was significantly larger in adipogenic cells (*p <* 0.01, Wilcoxon Rank Sum test). **(D)** We performed differential expression to identify genes with significant Cell Type:Reprogramming interactions. Qualitative analysis of gene expression in the latent space reveals cell type specific effects changes across the reprogramming trajectories. **(E)** We used Gene Ontology analysis to investigate the most significant Cell Type:Reprogramming interactions. Epithelial-to-mesenchymal transition (EMT) and extracellular matrix (ECM) gene sets were less downregulated in adipogenic cells after reprogramming (positive Adipogenic:Reprogramming interaction coefficients), while glycolytic gene sets were less upregulated (negative interaction coefficients). **(F)** Clustering a set of genes with significant Cell Type:Reprogramming interactions reveals prominent differences in the effect size of reprogramming across cell types. By contrast, effect sizes across age within the same cell type are fairly consistent.

**Figure S7:**
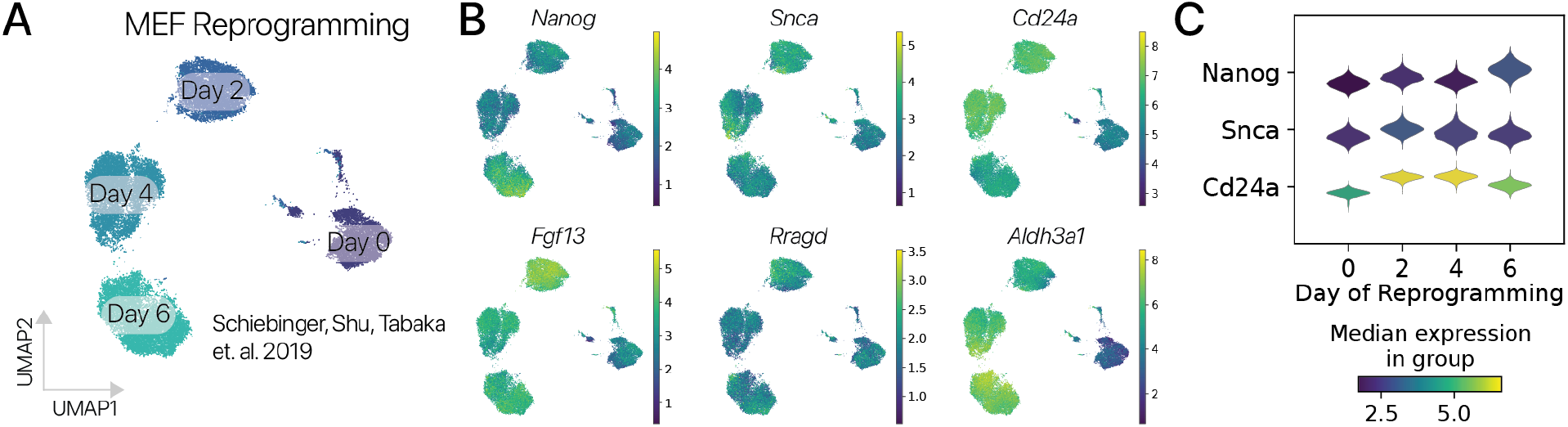
Pluripotency genes are activated in a subset of cells within 2-4 days of OSKM expression in mouse embryonic fibroblasts (MEFs). **(A)** UMAP projection of single cell mRNA profiles from MEFs undergoing reprogramming using a germline tetracycline-inducible OSKM allele [27]. Reprogramming induces novel cell states after only 2 days, consistent with strong effects we observe after a short OSKM pulse. **(B)** Expression of pluripotency markers (*Nanog, Rragd*) and early reprogramming response genes (others) across the MEF reprogramming timecourse. We observe induction of early reprogramming response genes consistent with our transient reprogramming data. We also observe activation of the *Nanog* pluripotency regulatory early in the timecourse (Day 2, 4), consistent with our transient reprogramming results. **(C)** Expression of pluripotency and early reprogramming genes across the MEF reprogramming timecourse. *Nanog* is significantly upregulated after only 2 days of reprogramming (*q <* 0.1, Monte Carlo estimation of log-fold change).

**Figure S8:**
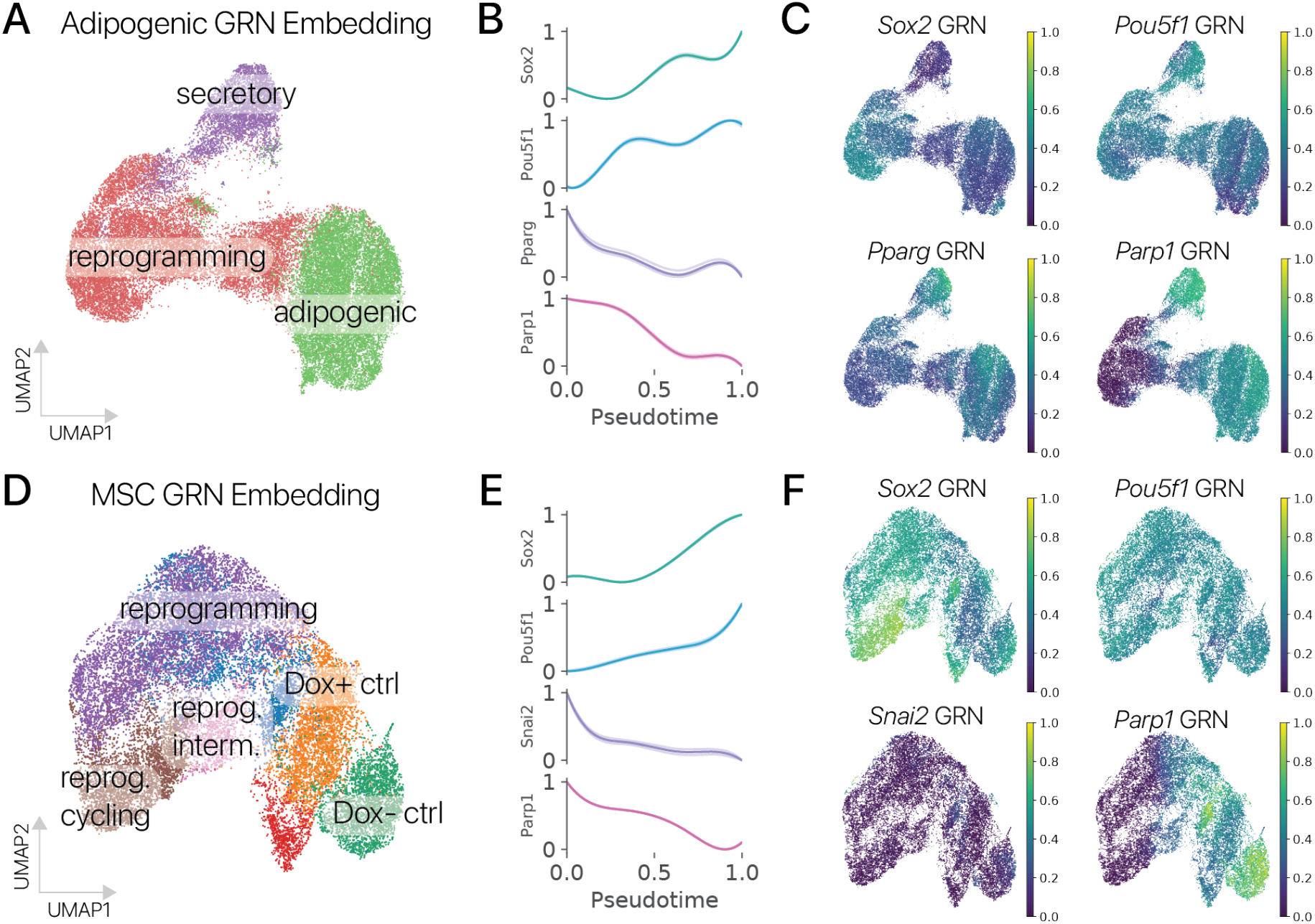
Gene regulatory network (GRN) analysis reveals the suppression of somatic cell identity GRNs during transient reprogramming. We extracted a directed GRN graph from the TRRUST database, linking murine transcription factors to direct target genes [30]. We then scored GRN activity in transiently reprogrammed adipogenic cells and MSCs using a rank based integration approach, adopted from SCENIC [29]. **(A)** Adipogenic cells embedded in a transcriptional space derived from GRN activity scores. GRN scores accurately capture the effects of reprogramming and cell state differences within control populations. **(B)** GRN scores across reprogramming pseudotime (higher is more reprogrammed). We found that pluripotency GRNs (*Sox2, Pou5f1*) increase as expected with reprogramming, while adipogenic identity GRNs (*Pparg, Parp1*) are suppressed (Wald test, Binomial GLM, *p <* 0.01). **(C)** Pluripotency GRNs (top) and adipogenic identity GRNs (bottom) projected in the GRN score latent space clearly distinguish control and transiently reprogrammed cells. **(D)** Muscle derived MSCs embedded using GRN activity scores. GRN scores again capture cell state changes induced by transient reprogramming. **(E)** Pluripotency GRNs increase as expected with reprogramming in MSCs, while mesenchymal identity GRNs (*Snai2, Parp1*) are suppressed (Wald test, Binomial GLM, *p <* 0.01). **(F)** Pluripotency (top) and mesenchymal identity (bottom) GRN scores in the GRN latent space show clear enrichment for reprogrammed and control populations respectively.

**Figure S9:**
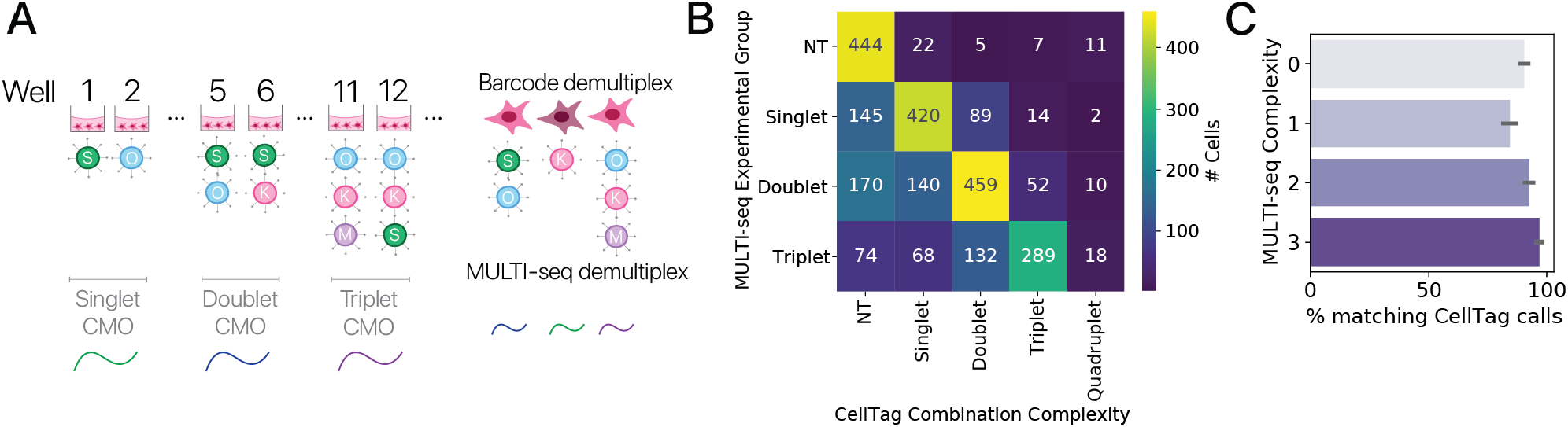
MULTI-seq labeling of combinatorial complexity and expressed barcode based demultiplexing show strong correspondence. **(A)** Experimental schematic. We performed a pilot pooled screen in an arrayed format and labeled the number of unique factors in each treatment condition using MULTI-seq. We then compared the MULTI-seq derived labels to our expressed barcode (CellTag) system for demultiplixing combinatorial perturbations. MULTI-seq label assignments are imperfect and expressed barcodes may suffer detection failures, so we do not expect perfect 1:1 correspondence. **(B)** Heatmap of correspondence between complexity labels derived from MULTI-seq and expressed barcodes for cells with a confident MULTI-seq derived label. We found that the expected complexity based on MULTI-seq barcodes is the dominant mode for expressed barcode classifications. We expect some elements in the lower triangle of the matrix due to (1) failed barcode detections or (2) cells that received less than the target number of unique factors in a given condition due to stochasticity in the transduction. We therefore consider all labels that are in the lower triangular matrix to be accurate. Very few cells fall outside the lower triangle, likely due to improper MULTI-seq barcode calls. **(C)** Correspondence of labels quantified across MULTI-seq treatment condition labels (mean + 95% CI). All complexities show high correspondence (>80% matching calls).

**Figure S10:**
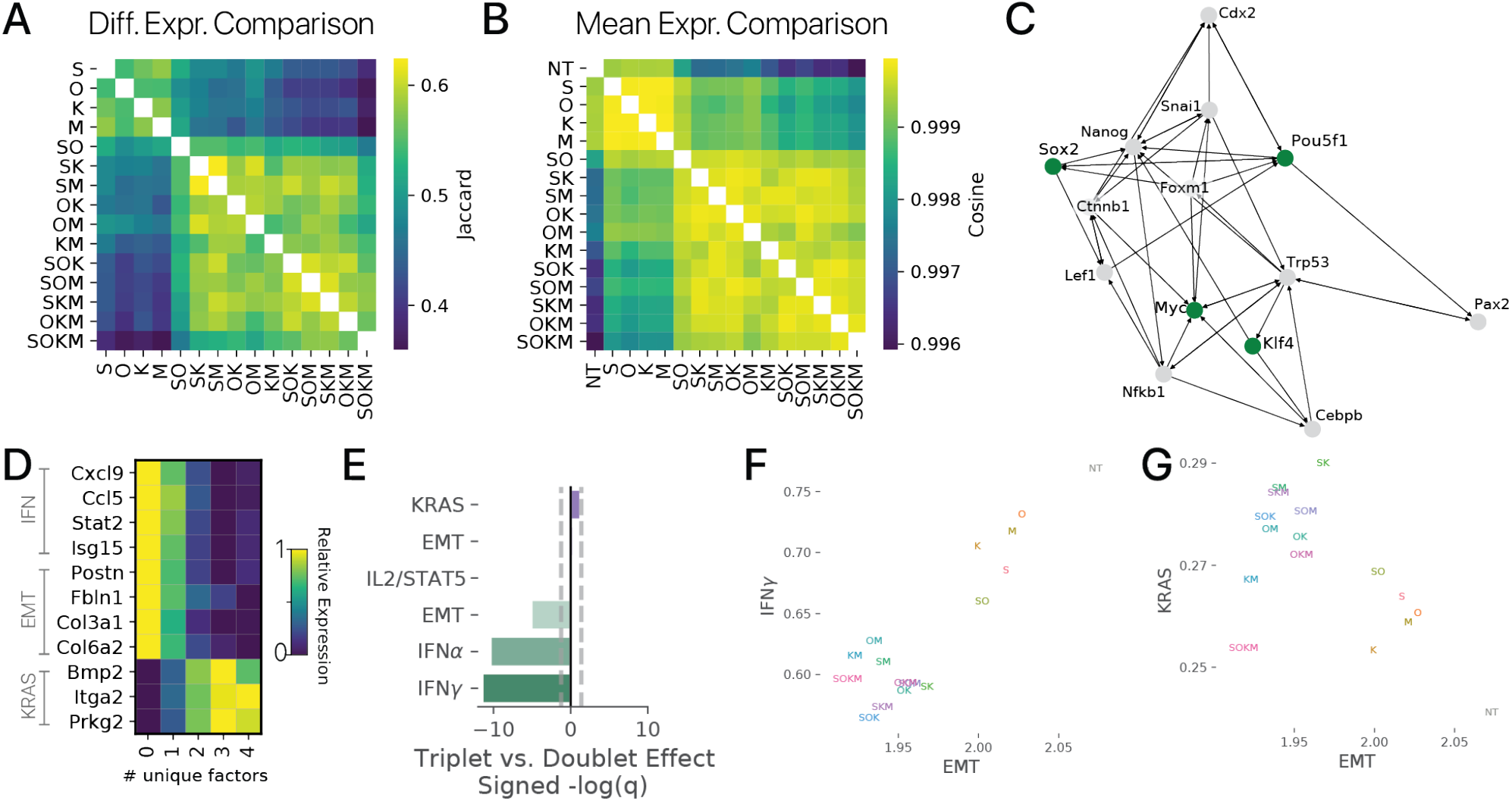
Higher-order combinations of Yamanaka factors exert similar transcriptional effects. **(A)** Comparison of differentially expressed gene (DEG) sets relative to the control for each perturbation. The Jaccard index measures the number of shared genes across two sets as a fraction of the total number of unique genes in both sets. Similar to results extracted from our perturbation classifier, we find that combinations of Yamanaka factors with similar complexity exert similar transcriptional effects after a transient pulse. **(B)** Comparison of the mean gene expression vectors for each Yamanaka Factor combination (NT: Not Treated controls) using the cosine similarity (positive values are more similar). We again found that higher order combinations exerted similar transcriptional effects. **(C)** Force directed layout visualization of connections within the pluripotency gene regulatory network extracted from the TRRUST database. Yamanka Factors are highlighted in green. All three factor subsets of the Yamanaka Factors have the potential to activate the fourth factor. **(D)** Comparison of interferon (IFN), epithelial-to-mesenchymal transition (EMT), and KRAS gene signatures across cells with different numbers of unique reprogramming factors. The largest differences between three and two factor combinations are stronger suppression of IFN and EMT programs and stronger activation of the KRAS signaling program. **(E)** Gene Ontology enrichment analysis with the MSigDB Hallmark gene sets confirms the gene level results in **(D). (F)** EMT and IFN*γ* program activity are tightly correlated (*r >* 0.9), suggesting that they are suppressed by a common mechanism. **(G)** EMT and KRAS program activities are anticorrelated (*r < −* 0.6), suggesting these effects likewise might share a mechanism during the reprogramming process.

**Figure S11:**
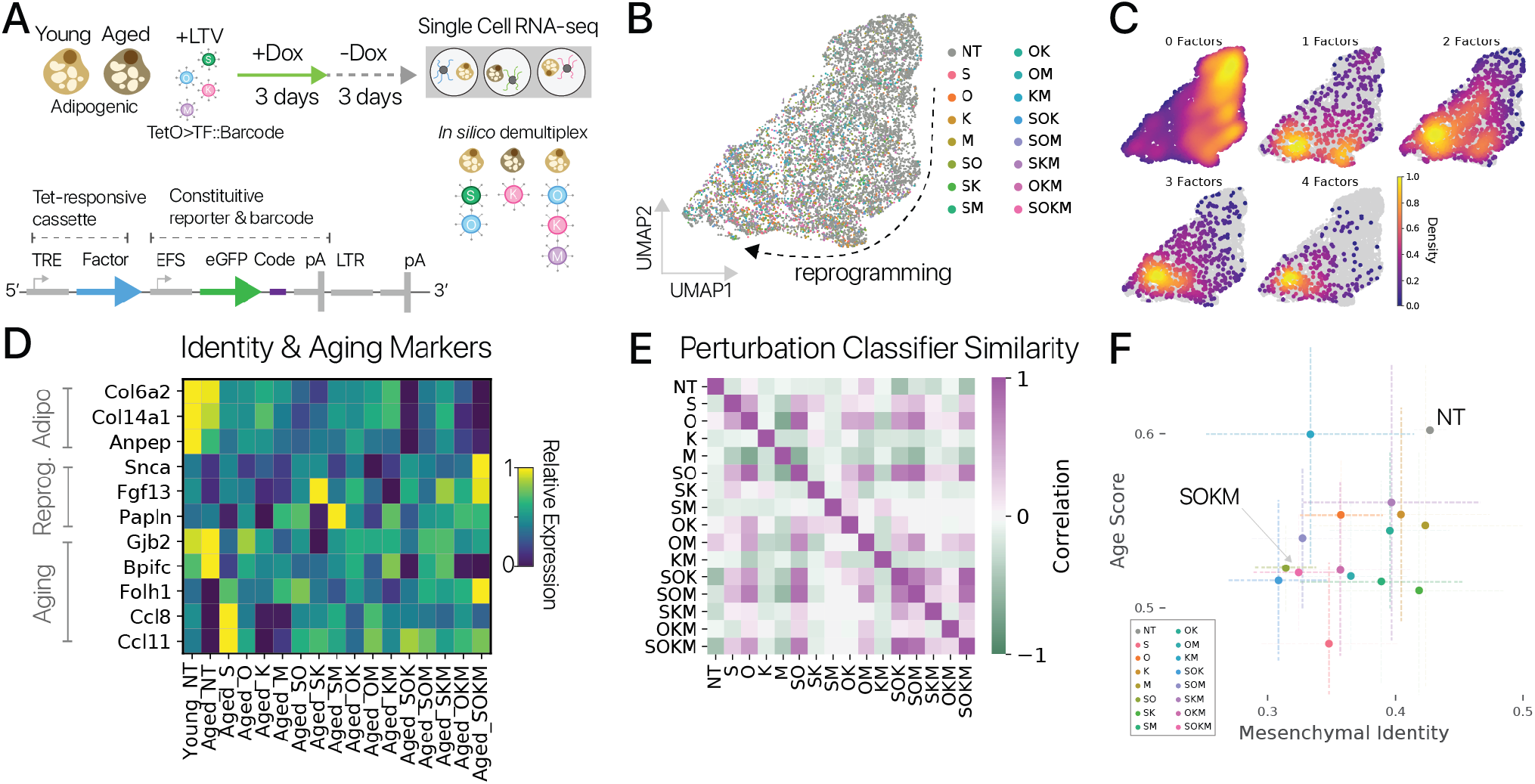
Pooled screening in adipogenic cells provides similar results to screening in MSCs. **(A)** Diagram of Yamanaka Factor pooled screening experiments. Young and aged adipogenic cells were transduced with lentiviruses each harboring a one inducible Yamanaka Factor with expressed barcodes (lower). Reprogramming was induced for a 3 day pulse/3 day chase (*n* = 5 animals per age across two independent experiments). Cells were profiled by single cell RNA-seq and unique combinations of Yamanaka Factors were demultiplexed *in silico* based on expressed barcodes. **(B)** Cells from the pooled screen embedded using scNym, projected with UMAP, and labeled with the detected reprogramming factors (9,000+ cells). **(C)** Density of cells perturbed with different numbers of reprogramming factors in the UMAP embedding. Higher order combinations show a larger transcriptional shift relative to control cells. **(D)** Adipogenic marker genes (top) decrease and reprogramming marker genes (center) increase as the combinatorial complexity (number of unique factors) increases. Aging genes (lower) likewise appear closer to the youthful level with more complex perturbations. **(E)** Similarity matrix between different Yamanaka Factor combinations extracted from a cell perturbation classification model. Classifier similarity is less dramatic than in MSCs, but the strongest similarities are again observed for higher order combinations of the Yamanaka Factors. **(F)** Mesenchymal cell identity scores derived from scNym models and aging gene set scores in aged cells reprogrammed with different factor combinations (mean scores *±* 95% CI). Many Yamanaka Factor combinations significantly decreased both the mesenchymal cell identity score and age score relative to aged control cells (NT) (Wald tests, *p <* 0.05). However, age scores and identity scores were not well-correlated, suggesting that cell identity suppression and rejuvenation are not tightly coupled (*ρ* = 0.19, *p >* 0.48, Spearman).

**Figure S12:**
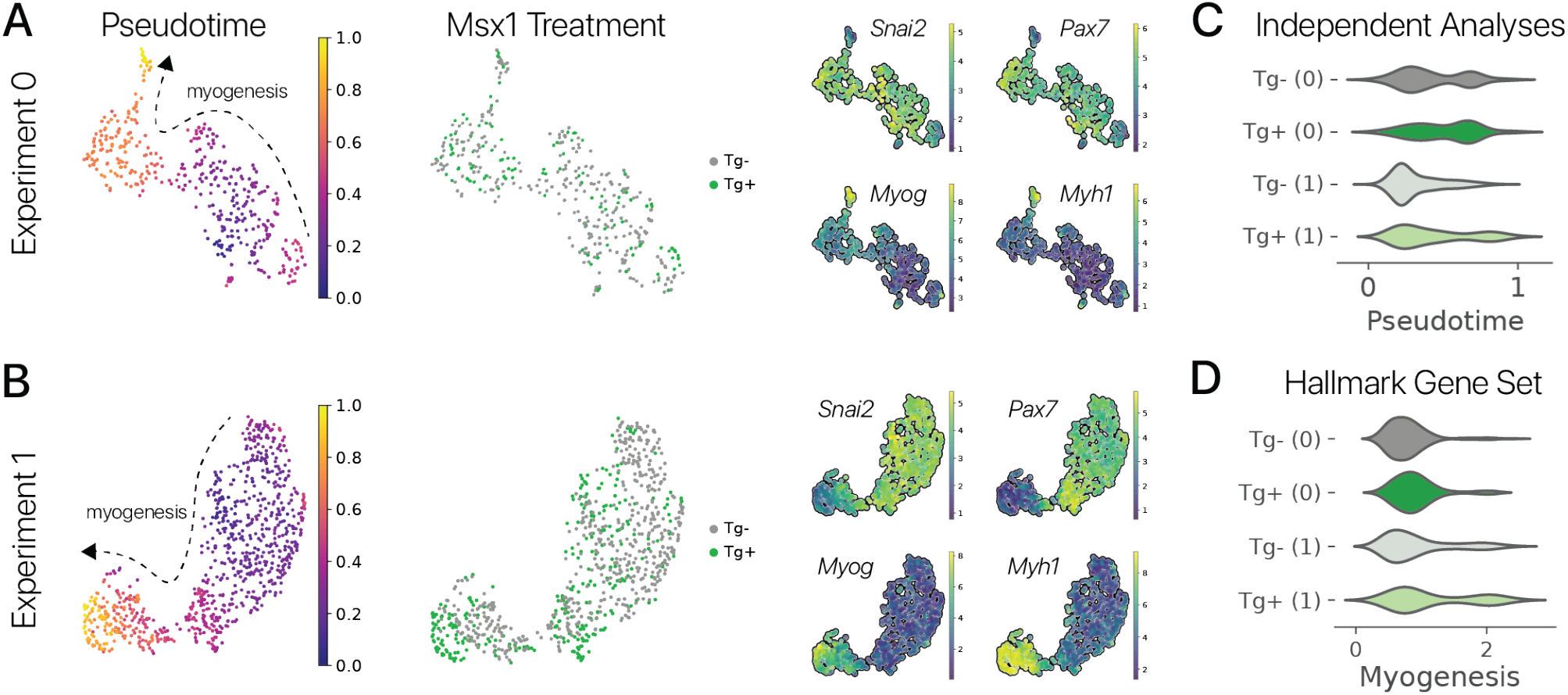
Pulsed reprogramming with the multipotency factor *Msx1* increases myogenesis in aged myogenic cells. Aged myogenic cells were treated with a transient pulse of *Msx1* in two separate experiments. Experiments were analyzed independently below. **(A)** Pseudotime analysis (left) and marker gene analysis (right) reveal a myogenic differentiation trajectory within both the first (*n* = 2 animals, 28 months old) and **(B)** second independent experiment (*n* = 3 animals, 30 months old). **(C)** We performed pseudotime analysis independently in each experimental condition. As in our integrated analysis (Fig. 4), transiently reprogrammed cells were significantly more differentiated than controls in both experiments (Wilcoxon Rank Sum test, *p <* 0.01). **(D)** We scored the activity of the Hallmark Myogenesis gene set (MSigDB). Both experiments showed significant increases in myogenic gene set activity after transient reprogramming treatment (Wilcoxon Rank Sum test, *p <* 0.01).

## Supplemental Tables

**Table S1:** Antibody information.

| Antigen | Supplier | Catalog |
| --- | --- | --- |
| CD31 | Biolegend | 102417 |
| CD45 | Biolegend | 103113 |
| ITGA7 | Thermo-Fischer | MA5-23555 |
| PDGFRα | Biolegend | 135916 |

